# Unmixing Spread Estimation Based on Residual Model in Spectral Flow Cytometry

**DOI:** 10.64898/2026.01.27.701929

**Authors:** Xiangming Cai, Sara Garcia-Garcia, Leo Kuhnen, Michaela Gianniou, Juan J. Garcia Vallejo

## Abstract

Advances in spectral flow cytometry have enabled the simultaneous measurement of dozens of markers across millions of cells within a single experiment. Despite the increasing maximum perplexity achievable in spectral panels, panel design remains constrained. A central obstacle is signal spread— unmixed fluorescence signal misattributed to unrelated channels—which reduces the resolution of cell populations. Here we introduce the Residual Model, a robust, scalable, and interpretable model-based approach for spread prediction during panel design. The Residual Model integrates statistical features derived from single-color controls and predicts spread under Ordinary Least Squares unmixing, the most widely used unmixing method. We demonstrate its reliable predictive performance across 141 single-color control samples measured on two instruments. To facilitate practical application, we developed the USERM R package, which implements the Residual Model and provides an out-of-box solution for interactive spread prediction and visualization.

## Introduction

Flow cytometry is one of the cornerstone techniques in life science research, enabling high-throughput characterization of cellular phenotypes^1^. Recent advances in spectral flow cytometry have further expanded these capabilities, allowing the simultaneous measurement of dozens of markers across millions of cells within a single experiment^2-8^. With continuous improvement, the panel complexity is rapidly increasing, with up to 50-plex panel already reported^9^.

However, designing a new panel remains a challenging task in practice^1,10^. A major obstacle is the signal spread, which arises during the unmixing process. The unmixing converts the raw detector-level measurement into analyzable marker intensities (for example, CD3). The signal spread was introduced as Spillover Spreading Error (SSE) in both conventional and spectral flow cytometry^11^, where the signal from one marker is partially misattributed to other markers^10,12^. This misattribution is derived by noise that perturbs the data, including both instrumental noise and physics noise (**Fig. 1A**). The physics noise reflects deviations of actual cellular emission from the averaged fluorescence signature. Spread increases background levels in negative populations, compromises resolution of cell populations at critical gating plots^13-15^, and ultimately constrains the design and performance of new panels.

**Figure 1.**
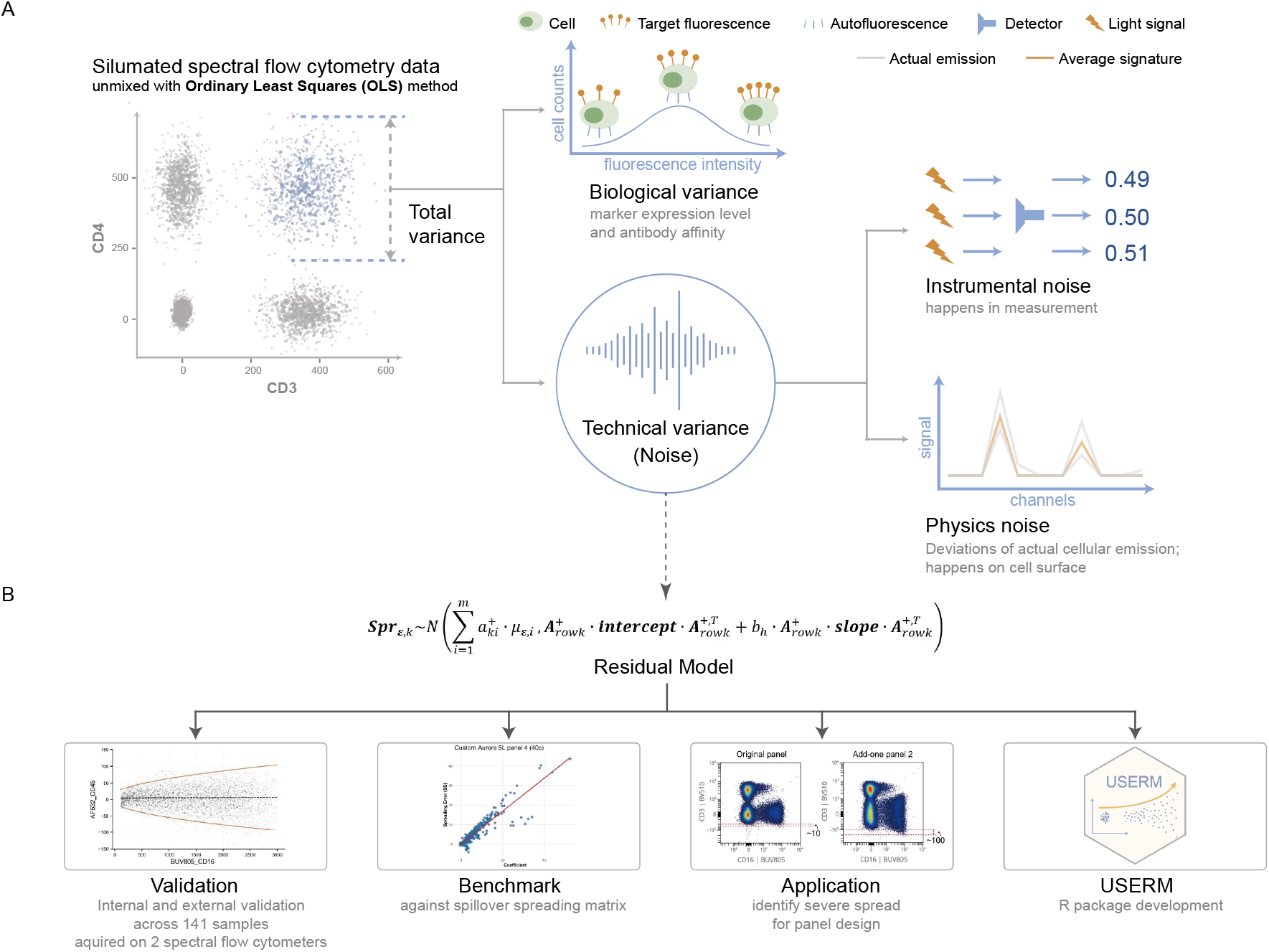
Spread component and research workflow. **(A)** Illustration of the total spread and its component structure. (B) Research workflow.

Therefore, it is critical to understand and predict spread at the early stages of spectral panel design. The Spillover Spreading Matrix (SSM) is the most widely used empirical tool for spread prediction^11^, which directly quantifies resolution across fluorescence channels based on measured and unmixed data. However, the SSM does not model the unmixing operation itself and therefore cannot explain the underlying causes of severe spread. Moreover, in practice, SSM is typically computed using bright single-color control (SCC) samples (such as CD4), which tend to exaggerate the level of spread observed in real experiments.

Although existing tools^16^ can provide estimation of spread, them offer limited insights into how spread happens, which could guide more effective solutions^11,17^. Most available tools attribute observed spread primarily to the overlap of fluorescence signatures. However, several additional factors substantially influence unmixed spread, including detector noise, inter-detector relationships, and fluorescence intensity (**Fig. 1B**). To address this gap, we developed Residual Model, which integrates these spread-related factors into a unified framework to model and predict spread under the Ordinary Least Squares (OLS) setting, the most widely used unmixing method in spectral flow cytometry.

The Residual Model boosts spectral flow cytometry panel design by enabling: (1) accurate prediction of spread for candidate panels, with flexibility to scale prediction according to expected fluorescence intensities; (2) interpretation and identification of the underlying sources of severe spread; (3) incorporation of custom SCC samples, which is particularly valuable when using new fluorophores, non-human peripheral blood mononuclear cell (PBMC) samples, or specialized experimental protocols that substantially alter fluorescence signature and spread; and (4) compatibility across different spectral flow cytometers, allowing SCC samples to be shared in a cross-instrument platform.

First, we characterize and formalize the spread phenomenon through the Residual Model and validate its predictive performance using real-world flow cytometry datasets. Second, we introduce and benchmark a practical tool, the Coefficient Matrix, which compactly summarizes and quantifies predicted spread. Finally, we develop the USERM R package, providing an out-of-box solution for researchers to access, apply and interactively visualize the Residual Model (**Fig. 1B**). We anticipate that this framework will deepen understanding of unmixed spread and substantially streamline spectral panel design by reducing the time and cost associated with iterative panel optimization.

## Results

### Residual Model for Unmixed Spread in Spectral Flow Cytometry

A major limitation of existing spread-prediction approaches is their inability to scale predicted spread as fluorescence intensity varies. To address this, the Residual Model integrates the fluorescence signature matrix, the variance-covariance structure of residual signals, and the intensity of SCC-specific fluorescence into a unified model. The model shows a fundamental linear relationship between the variance of spread and the SCC-specific fluorescence intensity. A detailed derivation is provided in the **Summplementary Methods**.

The aim of the Residual Model is to predict the spread of a SCC (stained with its SCC-specific fluorescence) when unmixed with a *n* -fluorescence signature matrix ( *n* >2). The signature matrix contains normalized signatures for the SCC-specific fluorescence, autofluorescence (AF) and ( *n* − 2) additional fluorescence. In this study, “spread” specificly refers to the unmixed signals appearing in these additional fluorescence. These spread signals originate from the residuals obtained when the raw data are unmixed using a minimal two- fluorescence signature matrix consisting only of the SCC-specific fluorescence and AF. By modelling these residual signals, the Residaul Model enables accurate prediction of spread when unmixing with a larger signature matrix.

### Reliable SCC Spread Prediction Enabled by Residual Model

To assess the performance of the Residual Model, we performed both internal and external validations using four SCC datasets generated on either the Xenith or Aurora 5L instruments: MC panel Cells Xenith (N = 31), MC panel Beads Xenith (N = 30), CD4 panel Xenith (N = 18), and CD4 panel Aurora (N = 62) (**Supplementary Fig. 1**). To apply the Residual Model, model parameters need to be estimated from SCC data. In internal validation, the SCC used for parameter estimation was identical to the SCC being predicted. In contrast, external validation involved estimating parameters from one SCC and predicting the spread of a different SCC (**Supplementary Fig. 1**). To ensure reliable assessment, only the positive population was included in the validation, which is characterized by stable morphology and sufficient cell counts. Model performance was quantified using the coverage rate, defined as the proportion of data points falling within the estimated 95% spread interval. A coverage rate approaching 0.95 indicates accurate prediction.

We first conducted internal validation using the MC panel Cells Xenith dataset (N = 31; **Fig. 2A**). Each SCC was unmixed with the Full MC panel signature matrix (31 colors + AF; **Supplementary Table 1**). Most SCC exhibited near-ideal coverage rates, ranging from 0.869 to 0.984. The only exception was PEVio770-TCRVb11, which showed lower coverage rates (0.728—0.956), attributable to the limited number of TCRVb11+ cells (∼456) in this SCC sample.

**Figure 2.**
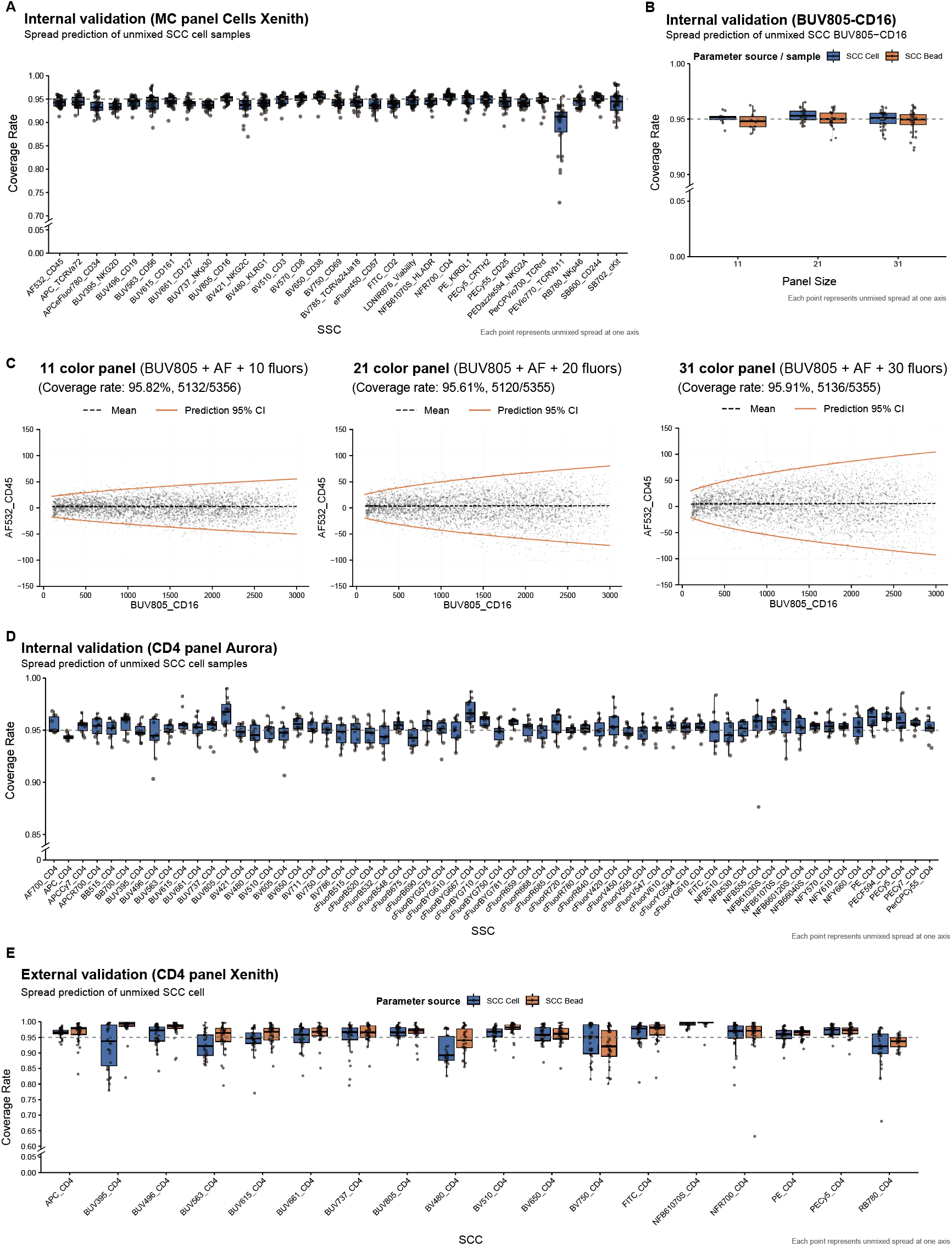
Validation of the predictive performance of Residual Model. **(A)** internal validation of the MC panel Cells Xenith dataset based on SCC cells. (B) internal validation of the SCC Cell BUV805-CD16. (C) internal validation of the SCC Cell BUV805-CD16 when unmixed with three panels of different panel size. (D) internal validation of the CD4 panel Aurora dataset based on SCC cells. (E) external validations of CD4 panel Xenith dataset.

Previous studies have shown that the fluorescence combination of the signature matrix significantly influences the spread signals^17^. To evaluate the robustness of the Residual Model under varying signature matrixes, we unmixed the SCC Cell BUV805-CD16 sample using two truncated MC panel signature matrixes (21 colors + AF; 11 colors + AF; **Fig. 2B; Supplementary Table 1**). Across these two and the Full MC panel configurations, the Residual Model yielded consistently high coverage rates along each negative axis (0.932 – 0.965). Of note, increased spread was observed when the signature matrix expands and the Residual Model accurately recapitulated this change (**Fig. 2C**).

We next evaluated model performance on SCC beads. SCC Bead BUV805-CD16 was unmixed with the same three MC panel signature matrixes (**Fig. 2B**), and the Residual Model again demonstrated strong performance, yielding satisfactory coverage rates across all axes.

Instrumental choice can also influence spread. To determine whether the Residual Model generates across platform, we performed internal validation using SCC cell data acquired on the Aurora 5L (CD4 panel Aurora; N = 62). For each SCC, 10 additional fluorescence were randomly selected and combined with the SCC-specific fluorescence and AF to construct a 12-fluorescence signature matrix. As shown in **Figure 2D**, ideal fits were achieved across all tested configurations.

To further evaluate generalizability of the Residual Model, we conducted external validation using the CD4 panel Xenith dataset (N = 18; **Supplementary Fig. 1**). Parameters were estimated from the MC panel Cells/Beads Xenith dataset and applied to predict the spread of the CD4 panel Xenith SCCs. Overall, most SCCs achieved coverage rates close to 0.95, indicating robust predictive performance. Predictions based on SCC cells and SCC beads were largely comparable. Representative cases with coverage rates below 0.8 or above 0.98 are shown in **Supplementary Figure 2**. Coverage rates below 0.8 were primarily attributable to a negatively predicted spread trend, likely resulting from suboptimal quality of the SCCs used for parameter estimation. In contrast, cases with coverage rates above 0.98 exhibited slightly exaggerated predictions, yet still captured the overall trend of the spread.

In summary, the Residual Model exhibits strong robustness across diverse experimental conditions. It reliably captures the spread of SCC irrespective of fluorescence composition, panel size, sample type (cells or beads), or instrument platform.

### Benchmarking the Residual Model Across Diverse Datasets

We benchmarked the performance of the Residual Model against SSM, the most commonly used approach for spread prediciton. In the Residual Model, a *Coefficient* is calculated to quantify the linear relationship between spread signal and the intensity of the SCC-specific fluroescence (**Fig. 3A**). Conceptually, coefficient is analogous to the spillover spreading (SS) index in the SSM; however, the coefficient is derived through a model-based estimation, whereas the SS is directly measured from unmixed SCC sample. Thus, a well-performed Residual Model should yield Coefficient that closely matchs the corresponding SS.

**Figure 3.**
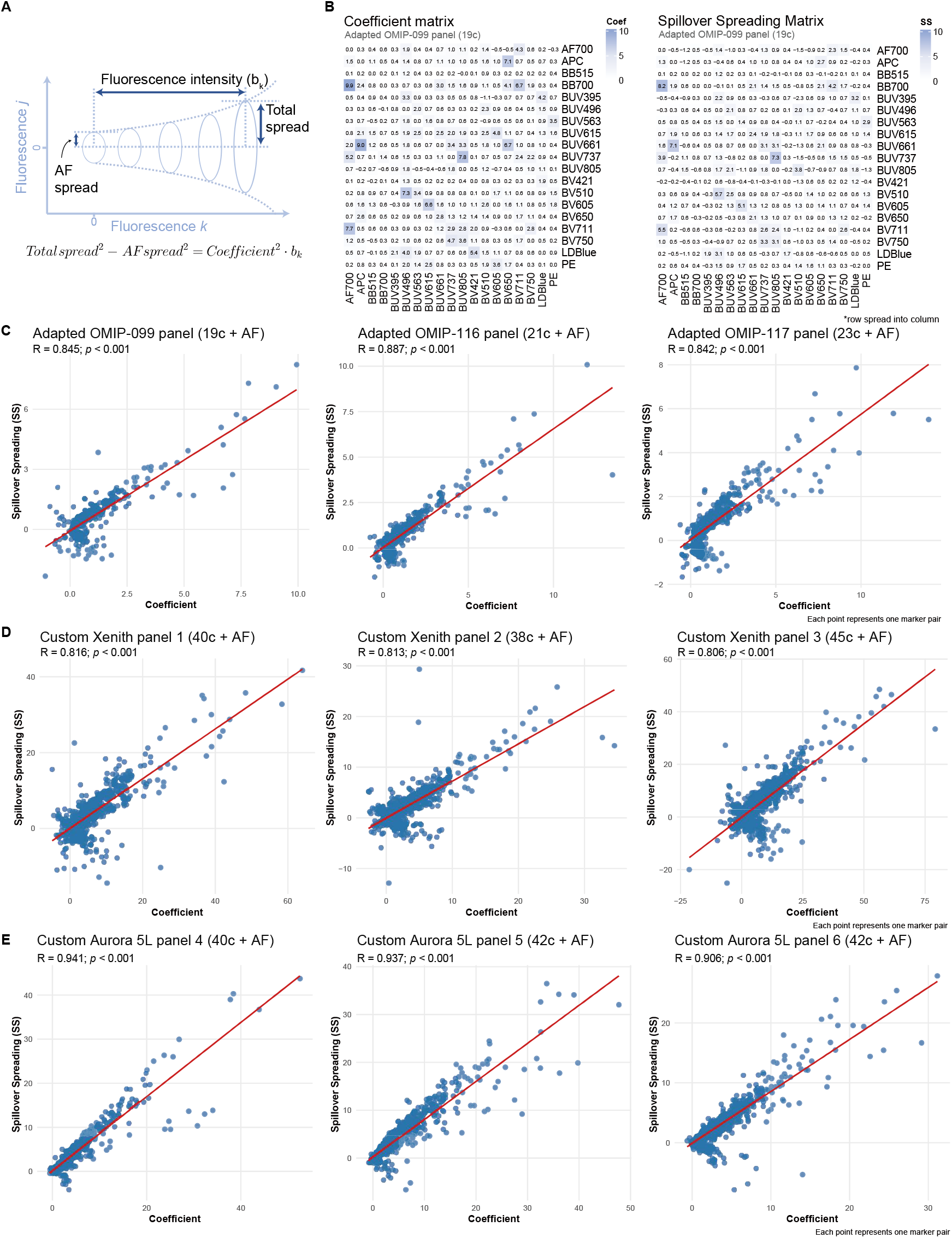
Benchmarking the Residual Model. **(A)** illustration of the Coefficient index. (B) Coefficient Matrix and SSM of adapted OMIP-099 panel. (C-E) Scatter plots of Coefficient versus SS in three adapted OMIP panels (C), three custom Xenith panels (D), and three custom Aurora 5L panel (E).

We first selected three OMIP panels that do not cross reference with each other (OMIP-099^18^, OMIP-116^19^, OMIP-117^20^). These panels are adapted based on fluorescence availability in our datasets (**Supplementary table 1)**. For each panel, we computed beth the SSM and a Coefficient Matrix with an identical structure. Representative matrixes for the adapted OMIP-099 panel is shown in **Figure 3B**. To quantitatively assess the correspondance between the Coefficient and the SS, we generated dot plots for the two indexes and conducted correlation analyses. All three adapted OMIP panels exhibited strong linear correlations (*p* < 0.001; **Fig. 3C**). Notably, severe spread found by the SSM can also be identified by the Coefficient Matrix. During computation, we observed negative SS and Coefficient values. Such value can arise in real datasets due to low positive events or stochastic variation. Existing tools may set negative SS to be zero for visualization purposes, which is accepatable because only substantial spread is of practical concern. Here, we retain the original values and most negative values were indeed small and close to zero.

To further challenge the Residual Model, we designed three large custom Xenith panel (∼40 colors), prioritizing Brilliant dyes, NovaFluor dyes, and cFluor dyes, respectively (**Supplementary table 1)**. Remarkably, the Coefficient and SS again demonstrated strong linear relationships (*p* < 0.001; **Fig. 3D**). To assess the performance across distict instruments, we additionally designed three large custom Aurora panel and observed similar consistent results (**Supplementary table 1; Fig. 3E**). Together, these analyses demonstrate that the Coefficient derived from the Residual Model reliably reflects the SS index and can identifies potential severe spread detected by SSM.

### Residual Model Identifies Severe Spread for Panel Design

Add a new fluorescence into a developing panel is a basic step in panel design. To assess whether the Residual Model can assist in this process, we analyzed a multicolor sample stained with the Full MC panel (31 colors; **Supplementary table 1**). The Coefficient Matrix generated for the Full MC panel served as the baseline reference (**Fig. 4A**). We then introduced an additional fluorescence whose cosine similarity to existing fluorescence was below 0.97, a commonly used threshold in panel design. A new Coefficient Matrix was computed to quantify the impact of adding new fluorescence on spread.

**Figure 4.**
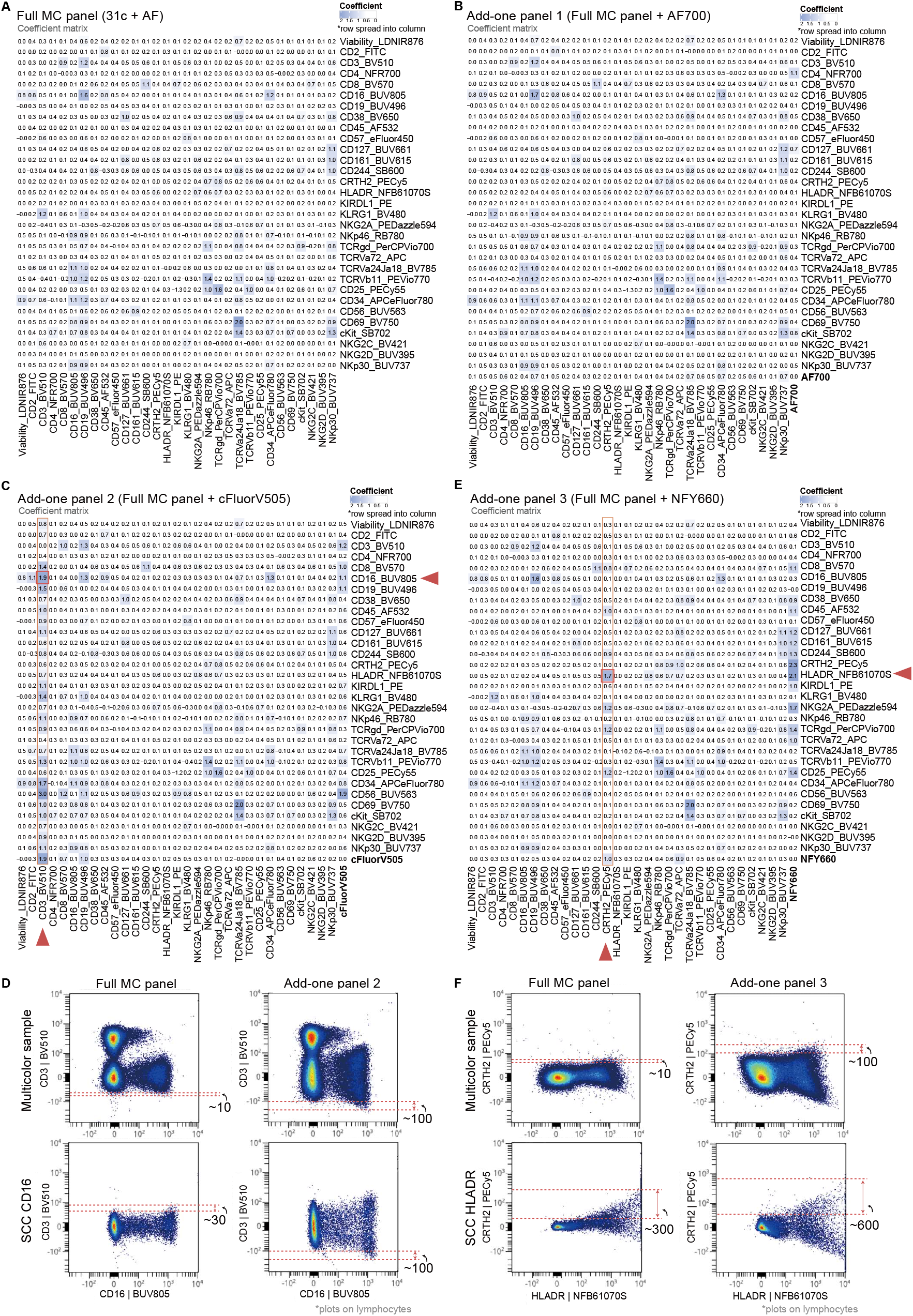
Residual Model Identifies Severe Spread for Panel Design. **(A)** Coefficient Matrix of Full MC panel. (B) Coefficient Matrix of Full MC panel + AF700. (C) Coefficient Matrix of Full MC panel + cFluorV505. (D) Scatter plots of CD16 versus CD3 when unmixed with Full MC panel or Full MC panel+ cFluorV505. (E) Coefficient Matrix of Full MC panel + NFY660. (F) Scatter plots of HLADR versus CRTH2 when unmixed with Full MC panel or Full MC panel+ NFY660.

Three representative fluorescence were tested. Incorporating of AF700 results in minimal changes to the spread (**Fig. 4B**), which was further supported by inspection of the N-by-N plots from the multicolor sample (data not shown). However, addition of cFluorV505 markedly increased the spread into CD3_BV510 (**Fig. 4C**), with the most pronounced contribution originating from CD16_BUV805. Scatter plots of CD3_BV510 versus CD16_BUV805 confirmed a substantial increase in background signals on the CD3_BV510 in the add-one panel (**Fig. 4D**). Also, the spread from CD16_BUV805 increased for around 3 times, compared with that in the Full MC panel. We also tested the NFY660, which led to increased spread into CRTH2_PECy5, accompanied by a clear rise in spread from HLADR_NFB61070S (**Fig. 4E**). This effect was similarly validated using scatter plots (**Fig. 4F**). Together, these results demonstrate that the Residual Model can identify fluorescence that introduce substantial spread, thereby enabling selection of fluorescence with minimal impact during panel design.

### USERM: Scalable Prediction Based on Residual Model

To facilitate the application of the Residual Model for researchers planning spectral flow cytometry experiments, we developed an R package named USERM (Unmixing Spread Estimation based on Residual Model) (**Fig. 5**). USERM provides a streamlined and user-friendly interface for implementing the Residual Model in routine panel design.

**Figure 5.**
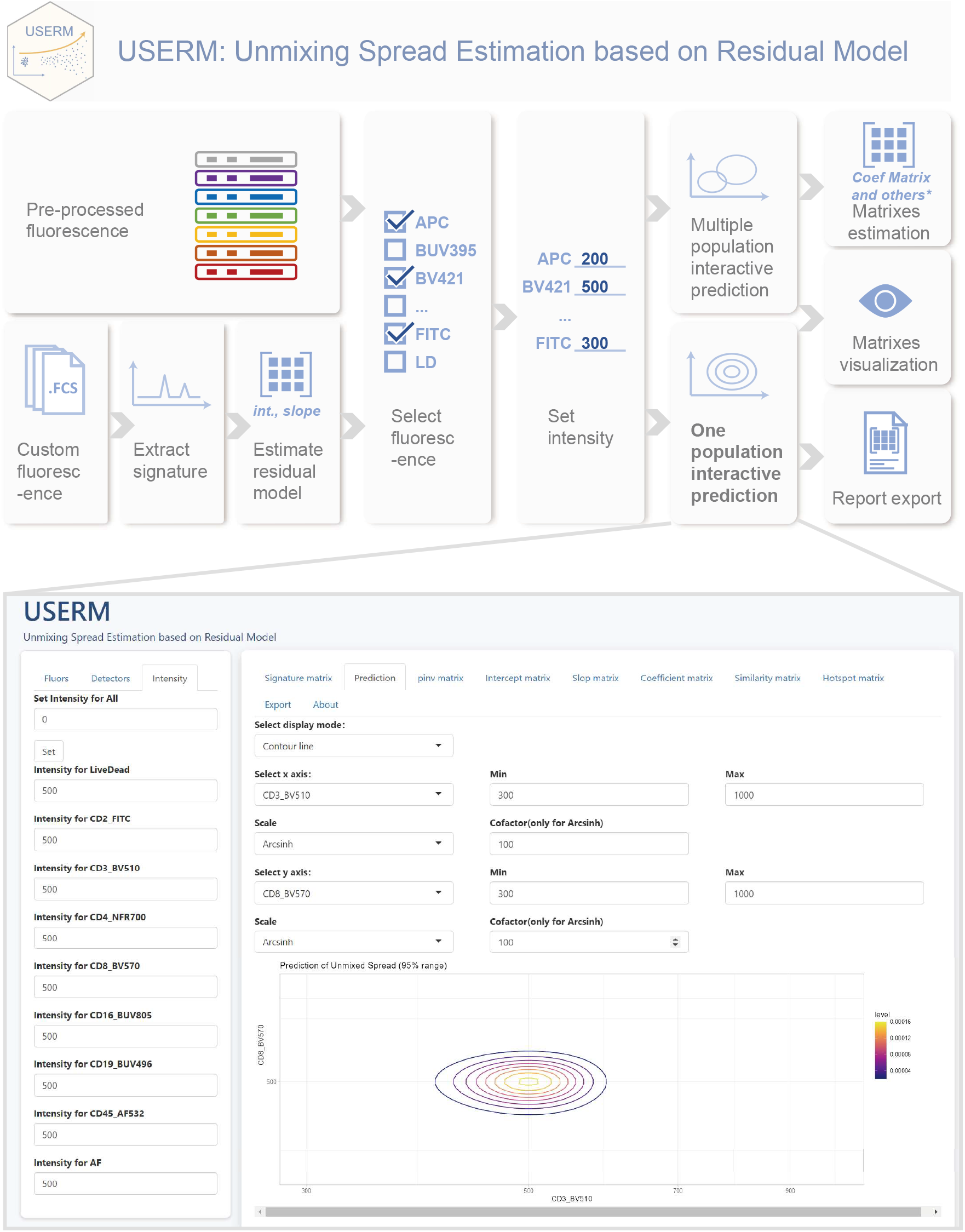
Illustration of USERM R package capability and workflow. ^*^other matrixes includes Spillover Spreading Matrix, Hotspot Matrix, and Cosine Similarity Matrix. The USERM provides an interactive interface for spread prediction, allowing users to adjust multiple parameters in real time.

To support immediate use, USERM incorporates preprocessed parameter sets derived from over 200 SCC samples acquired on Xenith and Aurora 5L instruments (**Supplementary Fig. 1**). These preprocessed data enable users to estimate spread without preparing SCC samples themselves. In addition to the built-in datasets, USERM supports custom SCC inputs, allowing users make spread prediction for panels containing alternative fluorescence or data generated on other spectral flow cytometry instruments. This flexibility ensures broad compatibility and promotes reproducible spread estimation across diverse experimental settings.

USERM provides flexible spread prediction functions for both single-population and multiple-population scenarios (**Fig. 5**). The package includes an interactive visualization function based on the *shiny* package. This function allows users to dynamically adjust fluorescence combination and intensities and intuitive explore their effects on predicted spread. What is more, USERM visualizes key components of the Residual Model, which allows interpretation of the underlying structure of the estimated spread and identification of the sources of severe spread.

USERM also supports the calculation and visualization of the Coefficient Matrix and other matrixes, including the SSM, Similarity Matrix, and Hotspot Matrix^17^. The Coefficient Matrix guides researchers to design a panel with acceptable spread, particularly at critical gating plot pairs where resolution is essential. Detailed usage instructions and vignettes are available in the USERM Github repository [https://github.com/xiangmingcai/USERM].

## Discussion

The spread issue is the key challenge in advancing spectral flow cytometry. In this study, we introduced the Residual Model, a model-based approach designed to quantitatively formalize and predict spread under the OLS setting. Through comprehensive internal and external validation, we demonstrated the robustness, applicability and generalizability of this approach. We further benchmarked its performance against SSM. To facilitate practical use, we developed the USERM R package, providing an out-of-box tool for panel design based on the Residual Model approach.

Traditionally, high expressed markers are paired with dim fluorescence, and vice versa, to achieve balanced unmixed signal. Here we propose an additional criterion: highly expressed markers should preferentially be paired with fluorescence that exhibit lower levels of spread within the panel. By following this criterion, researchers can achieve sufficient resolution to reliably distinguish cellular populations. The rationale is that even if a fluorescence has a strong tendency to generate spread, it will not substantially elevate background signal as long as the corresponding marker is expressed at low levels. In contrast, highly expressed markers (such as CD45, HLADR, and CD4) can cause considerable background signal even when the spread Coefficient or SS value is moderate or low. This also explains why we do not define a strict threshold for the Coefficient. Instead, decisions should be made by jointly considering both the spread potential and the expected fluorescence intensity of marker.

There are several limitations to this study. First, the Residual Model is derived based on the OLS unmixing method. For alternative unmixing algorithms^12^, further investigation will be required to adapt the model accordingly to accommodate these algorithms. Second, although the predictions are generally accurate, instances of suboptimal performance may occur. Reliable prediction depends on using high-quality SCC for parameter estimation. Specifically, high-quality should contain a sufficient number of cells whose fluorescence intensities span the expected range of the samples to be predicted. Otherwise, predicted results need to be interpreted with greater caution.

Another limitation is that, when applying the Residual Model for prediction, we assume that the AF of sample to be predicted is consistent with that of the sample used for parameter estimation. In practice, AF within a cell population can be complex and heterogeneous^21^. To assess the impact of this discrepancy, we estimated model parameters using SCC beads and applied them to predict spread in SCC cells (**Fig. 2E)**. The resulting coverage rates were comparable to those obtained based on SCC cells. This finding suggested that differences in AF between samples have only a limited effect on the prediction accuracy.

Finally, instrument setting, such as detector voltage, may vary slightly across institutions, potentially influencing observed spread patterns. However, when the vendor-recommended default settings are used, such variations are expected to have minimal impact. We recommend that users of USERM begin with our preprocessed fluorescence data. Alternatively, users may use SCC data generated within their own institutions if preferred.

The USERM provides a user-friendly access not only to the Residual Model approach, but also to general tools commonly used for panel design, including similarity matrix, hotspot matrix, and SSM. Current commercial panel-design platforms, such as Cytek^®^ Cloud, allow prediction only based on their own SCC datasets, which is acquired with their proprietary instruments. In contrast, USERM is released with more than 200 preprocessed SCC samples acquired on Xenith and Aurora 5L instruments. Users can expand this repository with SCC data generated in their own laboratories. This enables a single unified tool for panel design regardless of the instruments, sample types, species, or experiment protocol. Another advantage of USERM is that it is open-source, allowing the development of more advanced functionalities—for example, automated searches for alternative fluorescence.

To summarize, we developed the Residual Model as a scalable and interpretable model-based approach for spread prediction, offering practical support for panel design in spectral flow cytometry research. This approach has been implemented in the USERM R package, providing researchers with an out-of-box tool for spread prediction and comprehensive panel design.

## Method

### Materials and Data Acquisition

A total of four SCC datasets is generated in the study, including MC panel Cells Xenith (N = 31), MC panel Beads Xenith (N = 30), CD4 panel Xenith (complete; N = 73), and CD4 panel Aurora (N = 62) (**Supplementary Fig. 1**). PBMC from healthy donors were isolated from buffy coats (Sanquin Blood Bank, Amsterdam, The Netherlands) by density-gradient centrifugation using Lymphoprep (Serumwerk Bernburg, Germany). Cells were resuspended to 10-50×10^6^/ml in 20% DMSO in fetal bovine serum (FBS) and cryopreserved in liquid nitrogen until future use. Following thawing and cell recovery, cell viability was assessed prior to staining. The detailed staining protocol is provided in **Supplementary Methods**. Beads were also stained following the same procedure to generate the MC panel Beads Xenith dataset. Data acquisition was performed on either the Attune Xenith Flow Cytometer (Xenith) or the 5-laser Cytek^®^ Aurora (Aurora 5L) using default instrument settings. The MC panel datasets (X.C and L.K.) and CD4 panel datasets (M.G.) were prepared independently by different authors. A complete list of reagents and materials is provided in **Supplementary Materials**.

### Data preprocessing, unmixing, and analysis

Cleaning gates were used to isolate lymphocytes. First, singlets were identified by gating on FSC-A/FSC-H and SSC-A/SSC-H (using the 488-nm laser for Xenith). Subsequently, lymphocytes were geted based on FSC-A and SSC-A parameters. Negtative and positive populations were gated separately either using the peak detector channel (for validation in **Fig. 2**) or SCC-specific fluorescence after unmixing (for USERM use). Fluorescence signatures were extracted following the general principle^14^. Raw data were unmixed using the OLS algorithm. The Moore-Penrose pseudo-inverse matrix was computed to obtain the OLS solution. The pseudo-inverse was calculated using the *numpy*.*linalg*.*pinv* function from the *NumPy* library^22^ in Python or the *ginv* function from the *MASS* package in R. Parameters of the Residual Model were estimated for each SCC sample and subsequently used for prediction. Detailed scripts are available in the Github repository (https://github.com/xiangmingcai/USERM_paperMaterial).

### Residual Model architecture

The derivaiton of the Residual Model is base on three key empirical observations: (1) detector-specific residual signals follow a normal distribution; (2) residual signals across detectors correlate with each other; (3) the magnitude of spread is dependent on fluorescence intensity. These three empirical observations are examined and supported in the detailed derivation provided in the **Supplementary Materials**. We then demonstrate that, by characterizing the residual signals obtained when unmixing raw data using only SCC-specific fluorescence and AF, it becomes possible to predict the spread introduced by additional fluorescene when the signature matrix is expanded.

The unmixed spread signal for the *k*-th additional fluorescence can then be calculated using the following expression:

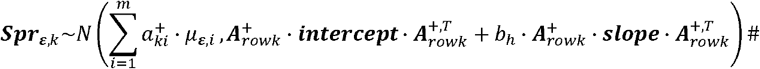

The ***A*** represents the signature matrix, with each column corresponding to the signature of a fluorescence. The 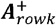 is the *k*-th row of the pseudo-inverse of the signature matrix. The 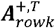 is the transpose of the 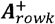. The *b*_*h*_ is the intensity of the SCC-specific fluorescence *h*. The Σ_*ε*_ denotes the variance-covariance matrix of the residual signals. By modelling the linear dependency between Σ_*ε*_ and *b*_*h*_, we obtain the corresponding ***intercept*** and ***slope*** matrixes that characterize this relationship. Detailed derivation steps can be found in the **Supplementary Method**.

This expression is referred to as the Residual Model. According to the Residual Model, the spread term follows a normal distribution and is linear related to the fluorescence intensity, with a coefficient given by 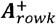 · **slope** · 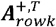 Note that to align with the definition of SS in SSM, the “Coefficient” reported in the Coefficient Matrix corresponds to the square root of 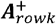 · **slope** · 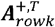.

### Benchmarking

We benchmarked the Residual Model-derived Coefficient against SS metric defined in the SSM. The Coefficient and SS share a similar interpretation, as both represent the square-root relationship between fluorescence intensity and spread. SSM calculation is implemented in the USERM R package via the *EstimateSSM* function, which follows the calculation procedures in the origianl publication. Positive and negative SCC populations are seperated in advance and stored within USERM. The *EstimateSSM* function imports these preprocessed data to compute the corresponding SSM. Pearson correlation was used to quantify the relationship between the Coefficent and SS.

### Development of USERM R package

The USERM R package is organized in a modular structure that provides core functions for SCC preprocessing, parameter estimation, spread prediction, and visualization. The package includes more than 200 preprocessed SCC samples to enable immediate use and reproducibility. It also supports user-provided SCC inputs to enable broader applicability across different experimental settings. USERM is implemented using established R libraries, including *flowCore, MASS, shiny, rmarkdown, DT, patchwork*, and *ggplot2*. Microsoft Copilot was used to assist in generating small code segments, all of which were carefully reviewed and verified by the authors (X.C.). Detailed documentations and vignettes are provided within the package (https://github.com/xiangmingcai/USERM).

## Data availability

All data needed to evaluate the conclusions in the paper are present in the paper and/or the Supplementary Materials. All datasets (MC panel Cells Xenith, MC panel Beads Xenith, CD4 panel Xenith, and CD4 panel Aurora) or are available in Figshare (link to Figshare).

## Code availability

Custom Python and R scripts used for analysis can be found in the Github repository (https://github.com/xiangmingcai/USERM_paperMaterial). The source code of USERM R package can be found on the GIthub repository (https://github.com/xiangmingcai/USERM).

## Acknowledgments

We thank Nick Rohrbacker, David Novo, and Pascutti Maria Fernanda for valuable conversations and insightful feedback. We also want to thank the MCCF (microscopy and cytometry core facility) of the Amsterdam UMC for their technical support in the laboratory.

This study was supported by the NICITA project (ADORE 2024-9-04) and China Scholarship Council (CSC; grant no. 202206090022).

## Author contributions

Conceptualization: X.C. Methodology: X.C. Investigation: X.C., S.G.G., J.J.G.-V. Data generation: X.C., L.K., M.G. Supervision: J.J.G.-V. Writing—original draft: X.C. Writing–review and editing: all authors. All authors contributed to the article and approved the submitted version. All authors had final responsibility for the decision to submit for publication.

## Figure legends

**Supplementary Figure 1 datasets generated in the study**.

**Supplementary Figure 2 representative good and poor prediction of external validation**.

(A) representative good prediction; (B) representative bad prediction with a coverage rate much higher than 0.95. (C) representative bad prediction with a coverage rate much lower than 0.95.

**Supplementary Fig3 the residual follows a normal distribution at each detector**.

(A-B) The distribution of residuals at each detector from the positive population of SCC cell BUV805-CD16(A) and FITC-CD2(B).

**Supplementary Fig4 The residuals at distinct detectors are correlated with each other**.

(A-B) The heatmap of correlation matrix of residuals at each detectors from the positive population (A) and negative population (B) of SCC cell BUV805-CD16 sample. (C-D) The heatmap of correlation matrix of residuals at each detectors from the positive population (C) and negative population (D) of SCC cell FITC-CD2 sample.

**Supplementary Fig5 The residuals are fluorescence intensity dependent**.

(A) example scatter plot shows the observed increase of spread along with the increase of fluorescence intensity. (B) Illustration of the subset method for the estimation of the observed increase of spread. (C) Example scatters of signature intensity (B) vs. coveriance between each detector and 781nm - 810/25-A (peak, as representation) from SCC cell BUV805-CD16 sample.

