## Supplementary Materials for "Unmixing Spread Estimation Based on Residual Model in Spectral Flow Cytometry"

### **Supplementary Methods**

#### 1 Construction and Derivation of Residual Model

The linear model is widely used to describe the mixing process:

$$\begin{aligned} \boldsymbol{Y}=\boldsymbol{A}\cdot\boldsymbol{B}+\boldsymbol{\varepsilon}\# \left( 0.1 \right) \end{aligned}$$

Here, $\boldsymbol{Y}$ denotes the detected raw signal matrix, where each column corresponds to an individual cell and each row to a detector. The $\boldsymbol{A}$ represents the signature matrix, with each column corresponding to the signature of a fluorescence. The $\boldsymbol{B}$ is the amount of fluorescence to be estimated. The $\boldsymbol{\varepsilon}$ accounts for the residual error of the formulation.

We first split the $\boldsymbol{B}$ in **Equation (0.1)** into two components:

$$\begin{aligned} \boldsymbol{B}=\boldsymbol{B}_{\boldsymbol{fluor}}+\boldsymbol{B}_{\boldsymbol{noise}}\# \left( 0.2 \right) \end{aligned}$$

Here, $\boldsymbol{B}_{\boldsymbol{fluor}}$ denotes the true amount of fluorescence, which is a fix value regardless of the signature matrix $\boldsymbol{A}$ used. The $\boldsymbol{B}_{\boldsymbol{noise}}$ represents the discrepancy between the estimated $\boldsymbol{B}$ and the true $\boldsymbol{B}_{\boldsymbol{fluor}}$.

Accordingly, we split $\boldsymbol{Y}$ into $\boldsymbol{Y}_{\boldsymbol{fluor}}$ and $\boldsymbol{Y}_{\boldsymbol{noise}}$:

$$\begin{aligned} \boldsymbol{Y}_{\boldsymbol{fluor}}=\boldsymbol{A}\cdot\boldsymbol{B}_{\boldsymbol{fluor}}\# \left( 0.3 \right) \end{aligned}$$

$$\begin{aligned} \boldsymbol{Y}_{\boldsymbol{noise}}=\boldsymbol{A}\cdot\boldsymbol{B}_{\boldsymbol{noise}}+\boldsymbol{\varepsilon}\# \left( 0.4 \right) \end{aligned}$$

The $\boldsymbol{Y}_{\boldsymbol{fluor}}$ represents the portion of raw signal that corresponds to the true fluorescence intensity $\boldsymbol{B}_{\boldsymbol{fluor}}$. It aligns precisely with a linear combination of average fluorescence signatures. So, there is no residual term for the $\boldsymbol{Y}_{\boldsymbol{fluor}}$**.** In contrast, the $\boldsymbol{Y}_{\boldsymbol{noise}}$ accounts for the noise part of the raw signal, which arises from instrumental noise or deviations between actual cellular emission and the average fluorescence signatures (**Fig. 1A**).

To improve clarity, we introduced a subscript for the $\boldsymbol{\varepsilon}$, defining $\boldsymbol{\varepsilon}_{\boldsymbol{noise}}\boldsymbol{=\varepsilon}$. Accordingly, **Equation (0.4**) can be rewritten as:

$$\begin{aligned} \boldsymbol{Y}_{\boldsymbol{noise}}=\boldsymbol{A}\cdot\boldsymbol{B}_{\boldsymbol{noise}}+\boldsymbol{\varepsilon}_{\boldsymbol{noise}}\# \left( 0.5 \right) \end{aligned}$$

The residual term $\boldsymbol{\varepsilon}_{\boldsymbol{noise}}$ is the portion of the noise signal that cannot be explained by the fluorescence signatures contained in $\boldsymbol{A}$.

**Equation (0.1)** can thus be reformulated as:

$$\begin{aligned} \boldsymbol{Y}=\boldsymbol{A}\cdot\boldsymbol{B}_{\boldsymbol{fluor}}+\boldsymbol{A}\cdot\boldsymbol{B}_{\boldsymbol{noise}}+\boldsymbol{\varepsilon}_{\boldsymbol{noise}}\# \left( 0.6 \right) \end{aligned}$$

In the ideal case where instrumental noise is absent and actual cellular emission perfectly conform to the fluorescence signatures, we have $\boldsymbol{Y}_{\boldsymbol{noise}}=\boldsymbol{0}$. Under this condition, the unmixed $\boldsymbol{B}$ values collapse onto a straight line in scatter plots, exhibiting no spread at all.

The OLS method is the most widely adopted method for estimating the intensity matrix $\boldsymbol{B}$. When $\boldsymbol{A}$ has full column rank, which means that the fluorescence signatures are linearly independent, the OLS solution can be obtained using the Moore-Penrose pseudo-inverse matrix $\boldsymbol{A}^{+}$. The estimation process can be expressed as:

$$\begin{aligned} \boldsymbol{A}^{+}\cdot\boldsymbol{Y}=\boldsymbol{A}^{+}\cdot\boldsymbol{A}\cdot\boldsymbol{B}_{\boldsymbol{fluor}}+\boldsymbol{A}^{+}\cdot\boldsymbol{A}\cdot\boldsymbol{B}_{\boldsymbol{noise}}+\boldsymbol{A}^{+}\cdot\boldsymbol{\varepsilon}_{\boldsymbol{noise}} \\ \boldsymbol{A}^{+}\cdot\boldsymbol{Y}=\boldsymbol{B}_{\boldsymbol{fluor}}+\boldsymbol{B}_{\boldsymbol{noise}}+\boldsymbol{A}^{+}\cdot\boldsymbol{\varepsilon}_{\boldsymbol{noise}}\#\left( 0.7 \right) \end{aligned}$$

Given that the $\boldsymbol{\varepsilon}_{\boldsymbol{noise}}$ is the residual component orthogonal to the column space of $\boldsymbol{A}$, it follows that:

$$\begin{aligned} \boldsymbol{A}^{+}\cdot\boldsymbol{\varepsilon}_{\boldsymbol{noise}}=\boldsymbol{0}\# \left( 0.8 \right) \end{aligned}$$

Substituting this into **Equation (0.7)** and multiplying both sides by $\boldsymbol{A}$, we obtain:

$$\begin{aligned} \boldsymbol{A}^{+}\cdot\boldsymbol{Y}=\boldsymbol{B}_{\boldsymbol{fluor}}+\boldsymbol{B}_{\boldsymbol{noise}} \\ \boldsymbol{A}\cdot\boldsymbol{A}^{+}\cdot\boldsymbol{Y}=\boldsymbol{A}\cdot\boldsymbol{B}_{\boldsymbol{fluor}}+\boldsymbol{A}\cdot\boldsymbol{B}_{\boldsymbol{noise}}\# \left( 0.9 \right) \end{aligned}$$

The difference between **Equation (0.6)** and (**0.9)** is:

$$\begin{aligned} \boldsymbol{Y-A}\cdot\boldsymbol{A}^{+}\cdot\boldsymbol{Y}=\boldsymbol{\varepsilon}_{\boldsymbol{noise}}\# \left( 0.10 \right) \end{aligned}$$

The **Equation (0.10)** illustrate how to calculate the residual signal of an OLS solution.

Now, considering two scenarios for spectral unmixing of a single-color control (SCC) sample labeled with CD3-FITC. In the first scenario, a 2-fluorescence signature matrix $\boldsymbol{A}_{\boldsymbol{2}}$ is used, which includes only FITC and autofluorescence (AF). In the second scenario, a bigger 32-fluorescence signature matrix $\boldsymbol{A}_{\boldsymbol{32}}$ is applied, which contains FITC, AF, and 30 additional fluorescence.

The raw signal $\boldsymbol{Y}$ can be expressed in both scenarios as:

$$\begin{aligned} \boldsymbol{Y}=\boldsymbol{A}_{\boldsymbol{2}}\cdot\boldsymbol{B}_{\boldsymbol{fluor,2}}+\boldsymbol{A}_{\boldsymbol{2}}\cdot\boldsymbol{B}_{\boldsymbol{noise,2}}+\boldsymbol{\varepsilon}_{\boldsymbol{noise,2}} \\ \boldsymbol{Y}=\boldsymbol{A}_{\boldsymbol{32}}\cdot\boldsymbol{B}_{\boldsymbol{fluor,32}}+\boldsymbol{A}_{\boldsymbol{32}}\cdot\boldsymbol{B}_{\boldsymbol{noise,32}}+\boldsymbol{\varepsilon}_{\boldsymbol{noise,32}} \end{aligned}$$

The $\boldsymbol{B}_{\boldsymbol{fluor,32}}$ is a $\left( \boldsymbol{32\times detectors} \right)$ matrix, and the $\boldsymbol{B}_{\boldsymbol{fluor,2}}$ is a $\left( \boldsymbol{2\times detectors} \right)$ matrix. By definition, $\boldsymbol{B}_{\boldsymbol{fluor}}$, the true amount of fluorescence remains consistent in both scenarios. So that, if $\boldsymbol{B}_{\boldsymbol{fluor,2}}=\left[ \begin{matrix} b_{\boldsymbol{fluor,2,FITC}} \\ b_{\boldsymbol{fluor,2,AF}} \end{matrix} \right]$, we have:

$$\begin{aligned} \boldsymbol{B}_{\boldsymbol{fluor,32}}\boldsymbol{=}\left[ \begin{matrix} \begin{matrix} b_{\boldsymbol{fluor,2,FITC}} \\ b_{\boldsymbol{fluor,2,AF}} \end{matrix} \\ 0 \\ \begin{matrix} \vdots\\ 0 \end{matrix} \end{matrix} \right] \\ \boldsymbol{A}_{\boldsymbol{2}}\cdot\boldsymbol{B}_{\boldsymbol{fluor,2}}=\boldsymbol{A}_{\boldsymbol{32}}\cdot\boldsymbol{B}_{\boldsymbol{fluor,32}} \end{aligned}$$

Consequently, the noise signal parts are also equivalent:

$$\begin{aligned} \boldsymbol{A}_{\boldsymbol{2}}\cdot\boldsymbol{B}_{\boldsymbol{noise,2}}+\boldsymbol{\varepsilon}_{\boldsymbol{noise,2}}=\boldsymbol{A}_{\boldsymbol{32}}\cdot\boldsymbol{B}_{\boldsymbol{noise,32}}+\boldsymbol{\varepsilon}_{\boldsymbol{noise,32}} \end{aligned}$$

Solving for $\boldsymbol{B}_{\boldsymbol{noise,32}}$ using the pseudo-inverse of $\boldsymbol{A}_{\boldsymbol{32}}$, we obtain:

$$\begin{aligned} \boldsymbol{B}_{\boldsymbol{noise,32}}=\boldsymbol{A}_{\boldsymbol{3}\boldsymbol{2}}^{\boldsymbol{+}}\cdot\boldsymbol{A}_{\boldsymbol{2}}\cdot\boldsymbol{B}_{\boldsymbol{noise,2}}\boldsymbol{+}\boldsymbol{A}_{\boldsymbol{3}\boldsymbol{2}}^{\boldsymbol{+}}\cdot\boldsymbol{\varepsilon}_{\boldsymbol{noise,2}}\boldsymbol{-}\boldsymbol{A}_{\boldsymbol{3}\boldsymbol{2}}^{\boldsymbol{+}}\cdot\boldsymbol{\varepsilon}_{\boldsymbol{noise,32}} \end{aligned}$$

Because of **Equation (0.8)**, $\boldsymbol{A}_{\boldsymbol{3}\boldsymbol{2}}^{\boldsymbol{+}}\cdot\boldsymbol{\varepsilon}_{\boldsymbol{noise,32}}\boldsymbol{=0}$. So that we have:

$$\begin{aligned} \boldsymbol{B}_{\boldsymbol{noise,32}}=\boldsymbol{A}_{\boldsymbol{3}\boldsymbol{2}}^{\boldsymbol{+}}\cdot\boldsymbol{A}_{\boldsymbol{2}}\cdot\boldsymbol{B}_{\boldsymbol{noise,2}}\boldsymbol{+}\boldsymbol{A}_{\boldsymbol{3}\boldsymbol{2}}^{\boldsymbol{+}}\cdot\boldsymbol{\varepsilon}_{\boldsymbol{noise,2}}\boldsymbol{\#} \left( 0.11 \right) \end{aligned}$$

The **Equation (0.11)** illustrate how the noise component estimated with a 32-fluorescece signature matrix can be derived from the estimation with a 2-fluorescence signature matrix and its residual. To further illustrate the derivation, we expand the relevant matrixes as follows.

Let $\boldsymbol{B}_{\boldsymbol{noise,2}}=\left[ \begin{matrix} b_{\boldsymbol{noise,2,FITC}} \\ b_{\boldsymbol{noise,2,AF}} \end{matrix} \right]$ represent the estimated intensity with a 2-fluorescence signature matrix $\boldsymbol{A}_{\boldsymbol{2}}$. When projected into the 32-fluroescence signature matrix space, the $\boldsymbol{A}_{\boldsymbol{3}\boldsymbol{2}}^{\boldsymbol{+}}\cdot\boldsymbol{A}_{\boldsymbol{2}}\cdot\boldsymbol{B}_{\boldsymbol{noise,2}}$ yields a vector in which the FITC and AF components are preserved, while the remaining 30 additional dimensions are zero:

$$\begin{aligned} \boldsymbol{A}_{\boldsymbol{3}\boldsymbol{2}}^{\boldsymbol{+}}\cdot\boldsymbol{A}_{\boldsymbol{2}}\cdot\boldsymbol{B}_{\boldsymbol{noise,2}}\boldsymbol{=}\left[ \begin{matrix} \begin{matrix} b_{\boldsymbol{noise,2,FITC}} \\ b_{\boldsymbol{noise,2,AF}} \end{matrix} \\ 0 \\ \begin{matrix} \vdots\\ 0 \end{matrix} \end{matrix} \right]\#\left( 0.12 \right) \end{aligned}$$

The full noise vector for the 32-fluorescence signature matrix $\boldsymbol{A}_{\boldsymbol{32}}$ is denoted as:

$$\begin{aligned} \boldsymbol{B}_{\boldsymbol{noise,32}}=\left[ \begin{matrix} \begin{matrix} b_{\boldsymbol{noise,32,FITC}} \\ b_{\boldsymbol{noise,32,AF}} \end{matrix} \\ b_{\boldsymbol{noise,32,add}\boldsymbol{1}} \\ \begin{matrix} \vdots\\ b_{\boldsymbol{noise,32,add}\boldsymbol{30}} \end{matrix} \end{matrix} \right]\#\left( 0.13 \right) \end{aligned}$$

The residual projection $\boldsymbol{A}_{\boldsymbol{3}\boldsymbol{2}}^{\boldsymbol{+}}\cdot\boldsymbol{\varepsilon}_{\boldsymbol{noise,2}}$ accounts for the discrepancy between $\boldsymbol{B}_{\boldsymbol{noise,32}}$ and $\boldsymbol{A}_{\boldsymbol{3}\boldsymbol{2}}^{\boldsymbol{+}}\cdot\boldsymbol{A}_{\boldsymbol{2}}\cdot\boldsymbol{B}_{\boldsymbol{noise,2}}$ in **Equation (0.12)** and (**0.13)**, and is given by:

$$\begin{aligned} \boldsymbol{A}_{\boldsymbol{3}\boldsymbol{2}}^{\boldsymbol{+}}\cdot\boldsymbol{\varepsilon}_{\boldsymbol{noise,2}}\boldsymbol{=}\left[ \begin{matrix} \begin{matrix} b_{\boldsymbol{noise,32,FITC}}\boldsymbol{-}b_{\boldsymbol{noise,2,FITC}} \\ b_{\boldsymbol{noise,32,AF}}\boldsymbol{-}b_{\boldsymbol{noise,2,AF}} \end{matrix} \\ b_{\boldsymbol{noise,32,add}\boldsymbol{1}} \\ \begin{matrix} \vdots\\ b_{\boldsymbol{noise,32,add}\boldsymbol{30}} \end{matrix} \end{matrix} \right] \end{aligned}$$

Generalizing this formulation to an arbitrary signature matrix $\boldsymbol{A}_{\boldsymbol{n}}$ with $\boldsymbol{n}$ fluorescence ($\boldsymbol{n>2}$ and $\boldsymbol{A}_{\boldsymbol{n}}$ includes fluorescence in $\boldsymbol{A}_{\boldsymbol{2}}$), we have:

$$\begin{aligned} \boldsymbol{A}_{\boldsymbol{n}}^{\boldsymbol{+}}\cdot\boldsymbol{A}_{\boldsymbol{2}}\cdot\boldsymbol{B}_{\boldsymbol{noise,2}}\boldsymbol{=}\left[ \begin{matrix} \begin{matrix} b_{\boldsymbol{noise,2,FITC}} \\ b_{\boldsymbol{noise,2,AF}} \end{matrix} \\ 0 \\ \begin{matrix} \vdots\\ 0 \end{matrix} \end{matrix} \right]\#(0.14) \\ \boldsymbol{B}_{\boldsymbol{noise,n}}=\left[ \begin{matrix} \begin{matrix} b_{\boldsymbol{noise,n,FITC}} \\ b_{\boldsymbol{noise,n,AF}} \end{matrix} \\ b_{\boldsymbol{noise,n,add}\boldsymbol{1}} \\ \begin{matrix} \vdots\\ b_{\boldsymbol{noise,n,add}\left( \boldsymbol{n-2} \right)} \end{matrix} \end{matrix} \right]\#(0.15) \\ \boldsymbol{A}_{\boldsymbol{n}}^{\boldsymbol{+}}\cdot\boldsymbol{\varepsilon}_{\boldsymbol{noise,2}}\boldsymbol{=}\left[ \begin{matrix} \begin{matrix} b_{\boldsymbol{noise,n,FITC}}\boldsymbol{-}b_{\boldsymbol{noise,2,FITC}} \\ b_{\boldsymbol{noise,n,AF}}\boldsymbol{-}b_{\boldsymbol{noise,2,AF}} \end{matrix} \\ b_{\boldsymbol{noise,n,add}\boldsymbol{1}} \\ \begin{matrix} \vdots\\ b_{\boldsymbol{noise,n,add}\left( \boldsymbol{n-2} \right)} \end{matrix} \end{matrix} \right]\#(0.16) \end{aligned}$$

For notational simplicity, we define this residual projection as:

$$\begin{aligned} \boldsymbol{Spr}_{\boldsymbol{\varepsilon}}\boldsymbol{=}\boldsymbol{A}_{\boldsymbol{n}}^{\boldsymbol{+}}\cdot\boldsymbol{\varepsilon}_{\boldsymbol{noise,2}}\# \left( 0.17 \right) \end{aligned}$$

From the **Equation (0.14)**, it is evident that the $\boldsymbol{A}_{\boldsymbol{2}}\cdot\boldsymbol{B}_{\boldsymbol{noise,2}}$ does not contribute to the additional dimensions present in $\boldsymbol{B}_{\boldsymbol{noise,}\boldsymbol{n}}$. Specifically, the components $b_{\boldsymbol{noise,n,add}\boldsymbol{1}}$,…, $b_{\boldsymbol{noise,n,add(n-2)}}$ arise solely from the unmixing of the residual $\boldsymbol{\varepsilon}_{\boldsymbol{noise,2}}$. This observation holds true for any signature matrix $\boldsymbol{A}_{\boldsymbol{n}}$ that includes FITC and AF in $\boldsymbol{A}_{\boldsymbol{2}}$.

To formalize this, we define the noise component attributable exclusively to the residual projection as:$\begin{aligned} \boldsymbol{B}_{\boldsymbol{noise,n,\varepsilon}}\boldsymbol{=}\left[ \begin{matrix} \begin{matrix} 0 \\ 0 \end{matrix} \\ b_{\boldsymbol{noise,n,add}\boldsymbol{1}} \\ \begin{matrix} \vdots\\ b_{\boldsymbol{noise,n,add}\left( \boldsymbol{n-2} \right)} \end{matrix} \end{matrix} \right]\# \left( 0.18 \right) \end{aligned}$

By construction, $\boldsymbol{B}_{\boldsymbol{noise,n,\varepsilon}}$ and $\boldsymbol{Spr}_{\boldsymbol{\varepsilon}}$ are equivalent with respect to the additional dimensions beyond FITC and AF. This component plays a critical role in characterizing the observed spread in unmixing, yet it remains challenging to estimate directly. In the present study, we address this challenge by modelling $\boldsymbol{Spr}_{\boldsymbol{\varepsilon}}$, thereby enabling accurate prediction of $\boldsymbol{B}_{\boldsymbol{noise,n,\varepsilon}}$.

1.1 **The detector-specific residual of noise signal** **follows a normal distribution**

To characterize the residual term $\boldsymbol{\varepsilon}_{\boldsymbol{noise,2}}$ when unmix with a 2-fluorescence signature matrix $\boldsymbol{A}_{\boldsymbol{2}}$, we need to understand its distribution across individual detector. Let

$$\begin{aligned} \boldsymbol{\varepsilon}_{\boldsymbol{noise,2}}\boldsymbol{=}\left[ \begin{matrix} \begin{matrix} s_{\boldsymbol{\varepsilon},1} \\ \vdots\end{matrix} \\ s_{\boldsymbol{\varepsilon},i} \\ \begin{matrix} \vdots\\ s_{\boldsymbol{\varepsilon},m} \end{matrix} \end{matrix} \right]\# \left( 1.1 \right) \end{aligned}$$

where $m$ represents the total number of detectors, and $s_{\boldsymbol{\varepsilon},i}$ denotes the residual of noise signal at the $i$-th detector.

To investigate the statistical properties of $s_{\boldsymbol{\varepsilon},i}$, we analyzed the positive populations of SCC samples. Each $s_{\boldsymbol{\varepsilon},i}$ was computed using **Equation (0.10)**. Empirical observations indicated that the distribution of $s_{\boldsymbol{\varepsilon},i}$ closely follows a normal distribution, as illustrated in **Supplementary Figure 3A**.

Accordingly, me model the detector-specific residual of noise signal as:

$$\begin{aligned} s_{\boldsymbol{\varepsilon},i}\boldsymbol{\sim}N\left( \mu_{s_{\boldsymbol{\varepsilon},i}}\text{ },\sigma_{s_{\boldsymbol{\varepsilon},i}}^{2} \right)\#(1.2) \end{aligned}$$

where $\mu_{s_{\boldsymbol{\varepsilon},i}}$ and $\sigma_{s_{\boldsymbol{\varepsilon},i}}^{2}$ denote the mean and variance of the noise distribution at detector $i$, respectively.

From **Supplementary Figure 3A**, it is obvious that although these $s_{\boldsymbol{\varepsilon},i}$ follows normal distribution centering 0, the residuals do not have constant variances. So, the homoscedasticity, one of the OLS assumptions, is violated. It is well-known that flow cytometry data do not meet the homoscedasticity assumption for OLS. However, it is common in the field to directly apply OLS method on flow cytometry data. To accurately model the unmixed spread with OLS method, we use specific measured $\mu_{s_{\boldsymbol{\varepsilon},i}}$ and $\sigma_{s_{\boldsymbol{\varepsilon},i}}^{2}$ in **Equation (1.2)** to describe the distribution of each $s_{\boldsymbol{\varepsilon},i}$, instead of using a consistent $N\left( 0\text{ },\sigma^{2} \right)$ following the OLS assumption.

1.2 **The** $s_{\boldsymbol{\varepsilon},i}$ **at each detector correlates with each other**

Now that we have the distribution of $\boldsymbol{\varepsilon}_{\boldsymbol{noise,2}}$, we can proceed to estimate the distribution of $\boldsymbol{Spr}_{\boldsymbol{\varepsilon}}$, as defined in **Equation (0.9)**. This begins with the following expansion:

$$\begin{aligned} \boldsymbol{Spr}_{\boldsymbol{\varepsilon}}=\boldsymbol{A}_{\boldsymbol{n}}^{\boldsymbol{+}}\cdot\boldsymbol{\varepsilon}_{\boldsymbol{noise,2}}=\boldsymbol{A}_{\boldsymbol{n}}^{\boldsymbol{+}}\cdot\left[ \begin{matrix} \begin{matrix} s_{\boldsymbol{\varepsilon},1} \\ \vdots\end{matrix} \\ s_{\boldsymbol{\varepsilon},i} \\ \begin{matrix} \vdots\\ s_{\boldsymbol{\varepsilon},m} \end{matrix} \end{matrix} \right]\#(2.1) \end{aligned}$$

If we write out $\boldsymbol{A}_{\boldsymbol{n}}^{\boldsymbol{+}}$, it can be expressed as:

$$\begin{aligned} \boldsymbol{A}_{\boldsymbol{n}}^{\boldsymbol{+}}=\left[ \begin{matrix} \begin{matrix} a_{11}^{+} & a_{12}^{+} \\ a_{21}^{+} & a_{22}^{+} \end{matrix} & \begin{matrix} \cdots& a_{1m}^{+} \\ \cdots& a_{2m}^{+} \end{matrix} \\ \begin{matrix} \vdots& \vdots\\ a_{n1}^{+} & a_{n2}^{+} \end{matrix} & \begin{matrix} \ddots& \vdots\\ \cdots& a_{nm}^{+} \end{matrix} \end{matrix} \right]=\left[ \begin{matrix} \begin{matrix} \boldsymbol{A}_{row1}^{+} \\ \vdots\end{matrix} \\ \boldsymbol{A}_{rowk}^{+} \\ \begin{matrix} \vdots\\ \boldsymbol{A}_{rown}^{+} \end{matrix} \end{matrix} \right]\#(2.2) \end{aligned}$$

Each row vector $\boldsymbol{A}_{rowk}^{+}$ is defined as:

$$\begin{aligned} \boldsymbol{A}_{rowk}^{+}=\left[ \begin{matrix} \begin{matrix} a_{k1}^{+} & a_{k2}^{+} \end{matrix} & \begin{matrix} \cdots& a_{km}^{+} \end{matrix} \end{matrix} \right]\#(2.3) \end{aligned}$$

Substituting **Equation (2.2)** into **Equation (2.1),** we obtain:

$$\begin{aligned} \boldsymbol{Spr}_{\boldsymbol{\varepsilon}}=\left[ \begin{matrix} \begin{matrix} \boldsymbol{A}_{row1}^{+} \\ \vdots\end{matrix} \\ \boldsymbol{A}_{rowk}^{+} \\ \begin{matrix} \vdots\\ \boldsymbol{A}_{rown}^{+} \end{matrix} \end{matrix} \right]\cdot\left[ \begin{matrix} \begin{matrix} s_{\boldsymbol{\varepsilon},1} \\ \vdots\end{matrix} \\ s_{\boldsymbol{\varepsilon},i} \\ \begin{matrix} \vdots\\ s_{\boldsymbol{\varepsilon},m} \end{matrix} \end{matrix} \right]\#(2.4) \end{aligned}$$

The resulting matrix $\boldsymbol{Spr}_{\boldsymbol{\varepsilon}}$ has dimensions $n \times1$, where $n$ corresponds to the number of fluorescence in $\boldsymbol{A}_{\boldsymbol{n}}$ and the single column represents the cell count. The $k$-th entry of $\boldsymbol{Spr}_{\boldsymbol{\varepsilon}}$ can be written as:

$$\begin{aligned} \boldsymbol{Spr}_{\boldsymbol{\varepsilon},k}=\boldsymbol{A}_{rowk}^{+}\cdot\left[ \begin{matrix} \begin{matrix} s_{\boldsymbol{\varepsilon},1} \\ \vdots\end{matrix} \\ s_{\boldsymbol{\varepsilon},i} \\ \begin{matrix} \vdots\\ s_{\boldsymbol{\varepsilon},m} \end{matrix} \end{matrix} \right] \\ =\left[ \begin{matrix} \begin{matrix} a_{k1}^{+} & a_{k2}^{+} \end{matrix} & \begin{matrix} \cdots& a_{km}^{+} \end{matrix} \end{matrix} \right]\cdot\left[ \begin{matrix} \begin{matrix} s_{\boldsymbol{\varepsilon},1} \\ \vdots\end{matrix} \\ s_{\boldsymbol{\varepsilon},i} \\ \begin{matrix} \vdots\\ s_{\boldsymbol{\varepsilon},m} \end{matrix} \end{matrix} \right] \\ =\sum_{i=1}^{m} a_{ki}^{+}\cdot s_{\boldsymbol{\varepsilon},i}\#(2.5) \end{aligned}$$

From **Equation (1.2)**, we know that each $s_{\boldsymbol{\varepsilon},i}$ follows a normal distribution. However, the weighted sum in **Equation (2.5)** requires estimation of the variance-covariance matrix of the vector $\boldsymbol{\varepsilon}_{\boldsymbol{noise,2}}$.

Using SCC samples, we observed that within the positive population, the detector specific $s_{\boldsymbol{\varepsilon},i}$ exhibit significant inter-detector correlation (**Supplementary Fig. 4**). Notably, this correlation is particularly pronounced between detector pairs that overlap with peak detectors in the signatures of positive fluorescence and AF.

Building on this observation, **Equation (2.5)** can be further refined to incorporate the joint distribution of the correlated $s_{\boldsymbol{\varepsilon},i}$ terms. Since each $s_{\boldsymbol{\varepsilon},i}$ follows a normal distribution and the detectors are statistically dependent, the propagated term $\boldsymbol{Spr}_{\boldsymbol{\varepsilon},k}$ can be modeled as a linear combination of correlated Gaussian variables:

$$\begin{aligned} \boldsymbol{Spr}_{\boldsymbol{\varepsilon},k}=\sum_{i=1}^{m} a_{ki}^{+}\cdot s_{\boldsymbol{\varepsilon},i} \\ \boldsymbol{\sim}\sum_{\boldsymbol{i}\boldsymbol{=}\boldsymbol{1}}^{\boldsymbol{m}} a_{ki}^{+}\cdot N\left( \mu_{\boldsymbol{\varepsilon},i},\sigma_{\boldsymbol{\varepsilon},i}^{2} \right) \\ \boldsymbol{\sim}N\left( \sum_{i=1}^{m} a_{ki}^{+}\cdot\mu_{\boldsymbol{\varepsilon},i},\boldsymbol{A}_{rowk}^{+}\cdot\text{Σ}_{\boldsymbol{\varepsilon}}\cdot\boldsymbol{A}_{rowk}^{+,T} \right)\#(2.6) \end{aligned}$$

Here, $\text{Σ}_{\boldsymbol{\varepsilon}}$denotes the variance-covariance matrix of the $\boldsymbol{\varepsilon}_{\boldsymbol{noise,2}}$:

$$\begin{aligned} \text{Σ}_{\boldsymbol{\varepsilon}}=\left[ \begin{matrix} \begin{matrix} Var\left( s_{\boldsymbol{\varepsilon},1} \right) & Cov\left( s_{\boldsymbol{\varepsilon},1} ,s_{\boldsymbol{\varepsilon},2} \right) \\ Cov\left( s_{\boldsymbol{\varepsilon},2} ,s_{\boldsymbol{\varepsilon},1} \right) & Var\left( s_{\boldsymbol{\varepsilon},2} \right) \end{matrix} & \begin{matrix} \cdots& Cov\left( s_{\boldsymbol{\varepsilon},1} ,s_{\boldsymbol{\varepsilon},m} \right) \\ \cdots& Cov\left( s_{\boldsymbol{\varepsilon},2} ,s_{\boldsymbol{\varepsilon},m} \right) \end{matrix} \\ \begin{matrix} \vdots& \vdots\\ Cov\left( s_{\boldsymbol{\varepsilon},m} ,s_{\boldsymbol{\varepsilon},1} \right) & Cov\left( s_{\boldsymbol{\varepsilon},m} ,s_{\boldsymbol{\varepsilon},2} \right) \end{matrix} & \begin{matrix} \ddots& \vdots\\ \cdots& Var\left( s_{\boldsymbol{\varepsilon},m} \right) \end{matrix} \end{matrix} \right]\#(2.7) \end{aligned}$$

and $\boldsymbol{A}_{rowk}^{+,T}$ represents the transpose of the row vector $\boldsymbol{A}_{rowk}^{+}$.

It is interesting to note that one of the OLS assumptions is that the residuals $s_{\boldsymbol{\varepsilon},i}$ are independent, which means the covariance between different residuals are expected to be 0. Apparently, this independence assumption of OLS is also violated when applying to real flow cytometry data. In this study, to accurately predict the spread in the OLS setting, we model with the measured variance-covariance matrix $\text{Σ}_{\boldsymbol{\varepsilon}}$, instead of using the independence assumption.

**1.3** $\boldsymbol{Spr}_{\boldsymbol{\varepsilon},k}$ **is fluorescence intensity dependent**

In SCC samples, we observed that the spread along the axis of positive fluorescence increases with fluorescence intensity (**Supplementary Fig. 5A**). This phenomenon was well-known by the field(5). To quantitatively assess this relationship, the positive population was stratified into 30 subsets based on fluorescence intensity level (**Supplementary Fig. 5B**). For each subset, we computed the corresponding $\text{Σ}_{\boldsymbol{\varepsilon}}$. Now, we use SCC for fluorescence $h$ as an example.

Analysis of individual entries in $\text{Σ}_{\boldsymbol{\varepsilon}}$, such as $Cov\left( s_{\boldsymbol{\varepsilon},i} ,s_{\boldsymbol{\varepsilon},j} \right)$, revealed that in majority cases, the covariance exhibits a linear relationship with fluorescence intensity (**Supplementary Fig. 5C**). By fitting each covariance term into a linear model, we obtained a pair of parameters—intercept and slope—for every detector pair ($i ,j$). This procedure yields two matrixes:

$$\begin{aligned} \boldsymbol{intercept} =\left[ \begin{matrix} \begin{matrix} {intercept}_{1,1} & {intercept}_{1,2} \\ {intercept}_{2,1} & {intercept}_{2,2} \end{matrix} & \begin{matrix} \cdots& {intercept}_{1,m} \\ \cdots& {intercept}_{2,m} \end{matrix} \\ \begin{matrix} \vdots& \vdots\\ {intercept}_{m,1} & {intercept}_{m,2} \end{matrix} & \begin{matrix} \ddots& \vdots\\ \cdots& {intercept}_{m,m} \end{matrix} \end{matrix} \right]\#(3.1) \end{aligned}$$

$$\begin{aligned} \boldsymbol{slope} =\left[ \begin{matrix} \begin{matrix} {slope}_{1,1} & {slope}_{1,2} \\ {slope}_{2,1} & {slope}_{2,2} \end{matrix} & \begin{matrix} \cdots& {slope}_{1,m} \\ \cdots& {slope}_{2,m} \end{matrix} \\ \begin{matrix} \vdots& \vdots\\ {slope}_{m,1} & {slope}_{m,2} \end{matrix} & \begin{matrix} \ddots& \vdots\\ \cdots& {slope}_{m,m} \end{matrix} \end{matrix} \right]\#(3.2) \end{aligned}$$

Using these matrixes, the covariance structure $\text{Σ}_{\boldsymbol{\varepsilon}}$ at a given fluorescence intensity $b_{h}$ can be estimated as:

$$\begin{aligned} \text{Σ}_{\boldsymbol{\varepsilon}}\left( b_{h} \right)=\boldsymbol{intercept}+b_{h}\cdot\boldsymbol{slope} \#(3.3) \end{aligned}$$

For a negative population, where $b_{k}=0$, this simplifies to:

$$\begin{aligned} \text{Σ}_{\boldsymbol{\varepsilon}}\left( b_{h} \right)=\boldsymbol{intercept=}\text{Σ}_{\boldsymbol{\varepsilon},AF}\#(3.4) \end{aligned}$$

It is obvious that for negative population, the signal only comes from the autofluorescence.

For positive population ($b_{h}>0$),

$$\begin{aligned} \text{Σ}_{\boldsymbol{\varepsilon}}\left( b_{h} \right)=\boldsymbol{intercept}+b_{h}\cdot\boldsymbol{slope} \\ =\text{Σ}_{\boldsymbol{\varepsilon},AF}+b_{h}\cdot\boldsymbol{slope}\#(3.4) \end{aligned}$$

Thus, the fluorescence-dependent change in covariance is given by:

$$\begin{aligned} \text{Σ}_{\boldsymbol{\varepsilon}}\left( b_{h} \right)-\text{Σ}_{\boldsymbol{\varepsilon},AF}=b_{h}\cdot\boldsymbol{slope} \#(3.5) \end{aligned}$$

Substituting this into **Equation (2.6)**, the distribution of $\boldsymbol{Spr}_{\boldsymbol{\varepsilon},k}$ becomes:

$$\begin{aligned} \boldsymbol{Spr}_{\boldsymbol{\varepsilon},k}\boldsymbol{\sim}N\left( \sum_{i=1}^{m} a_{ki}^{+}\cdot\mu_{\boldsymbol{\varepsilon},i},\boldsymbol{A}_{rowk}^{+}\cdot\boldsymbol{intercept}\cdot\boldsymbol{A}_{rowk}^{+,T}+b_{h}\cdot\boldsymbol{A}_{rowk}^{+}\cdot\boldsymbol{slope}\cdot\boldsymbol{A}_{rowk}^{+,T} \right)\#(3.6) \end{aligned}$$

From the **Equation (3.6)**, we concluded that in SCC data, the variance of $\boldsymbol{Spr}_{\boldsymbol{\varepsilon},k}$ —i.e., the spread along the negative fluorescence axis $k$— is linearly dependent on the fluorescence intensity $b_{h}$. The coefficient of this relationship, $\boldsymbol{A}_{rowk}^{+}\cdot\boldsymbol{slope}\cdot\boldsymbol{A}_{rowk}^{+,T}$, quantifies the rate at which spread increases with intensity. Meanwhile, $\boldsymbol{A}_{rowk}^{+}\cdot\boldsymbol{intercept}\cdot\boldsymbol{A}_{rowk}^{+,T}$ represents the baseline spread attributable to AF.

From here, we refer to **Equation (3.6)** as the **Residual Model**.

Again, the OLS expected that the residuals should be independent of the explanatory variable, $b_{h}$ in the case. However, this assumption is also violated. These violation of OLS assumptions when applying it to the real flow cytometry data emphasis the urgent need to model spread based on measured parameters, which is the purpose of this study, instead of simply accepting the assumption of OLS.

#### 2 spectral flow cytometry staining protocol

**Thawing**

1. Prewarm RPMI + 10%FCS at 37°C waterbath. When volume is big, warm at least for 30 min. Take out Live-Dead NIR876 from -20. Put in foil & drawer to protect from light.
2. Thaw cells in waterbath 37°C until there is a small clumb of ice visable.
3. Transfer cryo content to 15 ml tube.
4. Add 1 ml warm medium to the cryovial.
5. Slowly add 8 ml warm medium to the 15ml tube whilst gently rotating the tube.
6. Transfer the 1 ml from the cryo tube into the 15 ml tube.
7. Centrifuge 10 min, 1200 rpm.
8. Remove supernatant and resuspend pellet in 10ml PBS buffer.
9. Centrifuge 10 min, 1500 rpm @ 4°C.
10. Remove supernatant and resuspend pellet in 10ml FACS buffer. Take an aliquot for counting.
11. Centrifuge 10 min, 1500 rpm @ 4°C.
12. Dilute sample with **PBS** to **50 *10^6**/ml to easily transfer to staining plate. Put tube on ice.
13. Separate cells for LiveDead. Put half of the cells in 65 degree waterbath for 8mins. Cool down for 2-3mins, and mix with the other half of live cells.
14. Transfer cells to 96 V-bottom plate according to the plate outline; 100ul cells /well.
15. Add beads to 96 V-bottom plate according to the plate outline; 1 drop /well.

**Single color control Staining and multicolor/unstained test**

1. Prepare LD staining according to excel, **ABmix1**.
2. Centrifuge 5 min, 450g @ 4°C. Remove supernatant.
3. Add 150 µl prefix1 (diluted LD staining)/sample.  Incubate 30 min @ RT, in the dark.
4. Prepare surface staining according to excel, **ABmix2 and ABmix3**.
5. Add 150 µl PBS. Centrifuge 5 min, 450g @ 4°C. Remove supernatant.
6. Add 150 µl FCS buffer, centrifuge 5 min, 450 g @ 4°C. Remove supernatant
7. Add **ABmix2** mix/sample to the pellet and mix. Incubate 20 min @ RT, in the dark.
8. Add **ABmix3** mix/sample to the pellet and mix. Incubate 30 min @ RT, in the dark.
9. Centrifuge staining plate 450g for 5 minutes @ 4°C. Remove supernatant. Resuspend each well in 150 µl FACS buffer. (1st wash)
10. Centrifuge staining plate 450g for 5 minutes @ 4°C. Remove supernatant. Resuspend each well in 150 µl FACS buffer. (2nd wash)
11. Resuspend cells and beads in 150 ul of ice cold PBS and transfer to Eppendorf.
12. Fixation for cells and beads. Add 50 ul of 4% PFA while vortexing and keep on ice in the dark for 15 min.
13. Use plate centrifuge for beads. Use tube centrifuge for cells.
14. Centrifuge on table-top microcentrifuge for 10 seconds at 15000 rpm.
15. Remove 160 ul of supernatant. Add 160 ul of FCS buffer.
16. Centrifuge on table-top microcentrifuge for 10 seconds at 15000 rpm.
17. Remove 160 ul of supernatant. Add 160 ul of FCS buffer.
18. Centrifuge on table-top microcentrifuge for 10 seconds at 15000 rpm.
19. Remove 160 ul of supernatant. Add up to be **1000** ul/tube. Transfer to measuring tube.

Note: For multicolor sample, ABmix1 is diluted LD; ABmix2 is prestain mix of CD25, CD45, TCR γ/δ, TCR Vα24 Jα18, TCR Vβ11, TCR Vα7.2; ABmix3 is the mix of rest antibodies. For SCC, the same procedures are used with only one corresponding antibody used.

### **Material list**

| Reagent | Cat no. |
| --- | --- |
| BD Horizon™ Human CD4 Fluorochrome Evaluation Kit | 566352 |
| Invitrogen™ NovaFluor™ CD4 Label Characterization Kit 1 | 17216576 |
| Invitrogen™ NovaFluor™ CD4 Label Characterization Kit 2 | 17236576 |
| Cytek® cFluor® CD4 Kit for V, B, YG, R Lasers | R7-40003 |
| BD OptiBuild™ BUV395 Mouse Anti-Human CD314 (NKG2D) | 743561 |
| BD OptiBuild™ BUV496 Mouse Anti-Human CD19 | 741141 |
| BD Horizon™ BUV563 Mouse Anti-Human CD56 | 612928 |
| BD OptiBuild™ BUV615 Mouse Anti-Human CD161 | 751151 |
| BD OptiBuild™ BUV661 Mouse Anti-Human CD127 | 749981 |
| BD OptiBuild™ BUV737 Mouse Anti-Human CD337 (NKp30) | 749128 |
| CD16 Monoclonal Antibody (eBioCB16 (CB16)), Brilliant Ultra Violet™ 805, eBioscience™ | 368-0168-42 |
| BD OptiBuild™ BV421 Mouse Anti-Human CD159c (NKG2C) | 748169 |
| CD57 Monoclonal Antibody (TB01 (TBO1)), eFluor™ 450, eBioscience™ | 48-0577-42 |
| BD Horizon™ BV480 Mouse Anti-Human KLRG1 | 568862 |
| Brilliant Violet 510™ anti-human CD3 Antibody | 317332 |
| Brilliant Violet 570™ anti-human CD8a Antibody | 301038 |
| CD244 Monoclonal Antibody (eBioC1.7 (C1.7)), Super Bright™ 600, eBioscience™ | 63-5838-42 |
| Brilliant Violet 650™ anti-human CD38 Antibody | 356620 |
| CD117 (c-Kit) Monoclonal Antibody (104D2), eFluor™ 450, eBioscience™ | 67-1178-42 |
| BD Horizon™ BV750 Mouse Anti-Human CD69 | 568285 |
| Brilliant Violet 785™ anti-human TCR Vα24-Jα18 (iNKT cell) Antibody | 342932 |
| CD2 Monoclonal Antibody (RPA-2.10), FITC, eBioscience™ | 11-0029-42 |
| CD45 Monoclonal Antibody (HI30), Alexa Fluor™ 532, eBioscience™ | 58-0459-42 |
| HLA-DR Monoclonal Antibody (LN3), NovaFluor™ Blue 610-30S, eBioscience™ | H049T03B05 |
| TCRγ/δ Antibody, anti-human | 130-113-514 |
| BD Horizon™ RB780 Mouse Anti-Human CD335 (NKp46) | 569330 |
| CD158a Recombinant Rabbit Monoclonal Antibody (124), PE | MA5-40963 |
| PE/Dazzle™ 594 anti-human CD159a (NKG2A) Antibody | 375122 |
| CD294 (CRTH2) Monoclonal Antibody (BM16), PE-Cyanine5, eBioscience™ | 15-2949-42 |
| CD25 Monoclonal Antibody (BC96), PE-Cyanine5.5, eBioscience™ | 35-0259-42 |
| TCR Vβ11 Antibody, anti-human, REAfinity™ | 130-126-304 |
| APC anti-human TCR Vα7.2 Antibody | 351708 |
| CD4 Monoclonal Antibody (SK3 (SK-3)), NovaFluor™ Red 700, eBioscience™ | H001T03R03 |
| CD34 Monoclonal Antibody (4H11), APC-eFluor™ 780, eBioscience™ | 47-0349-42 |
| LIVE/DEAD™ Fixable Near IR (876) Viability Kit, for 808 nm excitation | L34981 |
| eBioscience Flow Cytometry Staining Buffer | 00-4222-26 |
| CellBlox Plus blocking buffer (use with NovaFluor Ab Conjugates) | C001T03F01 |
| Anti-Hu Fc Receptor Binding Inhibitor | 14-9161-73 |
| Briliant Staing Buffer | 00-4409-42 |
| UltraComp eBeads Plus Compensation Beads | 01-3333-42 |

### **Supplementary Tables**

#### **Supplementary Table 1 panels list**.

Panels used in the study.

See additional files

#### **Supplementary Table 2 Figure data**

Result data for Figure 2.

See additional files

### **Supplementary Figs**

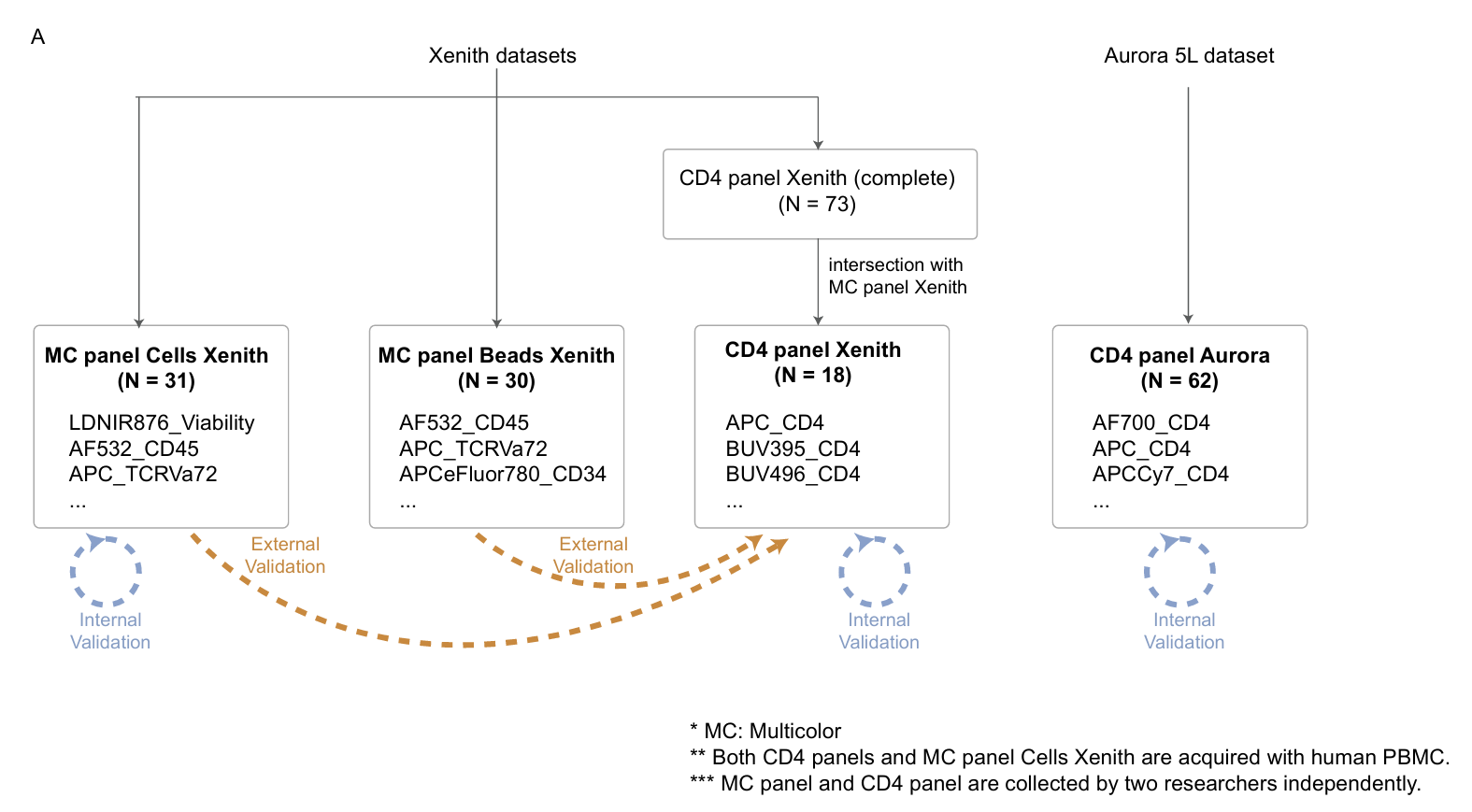

#### **Supplementary Figure 1 datasets generated in the study.**

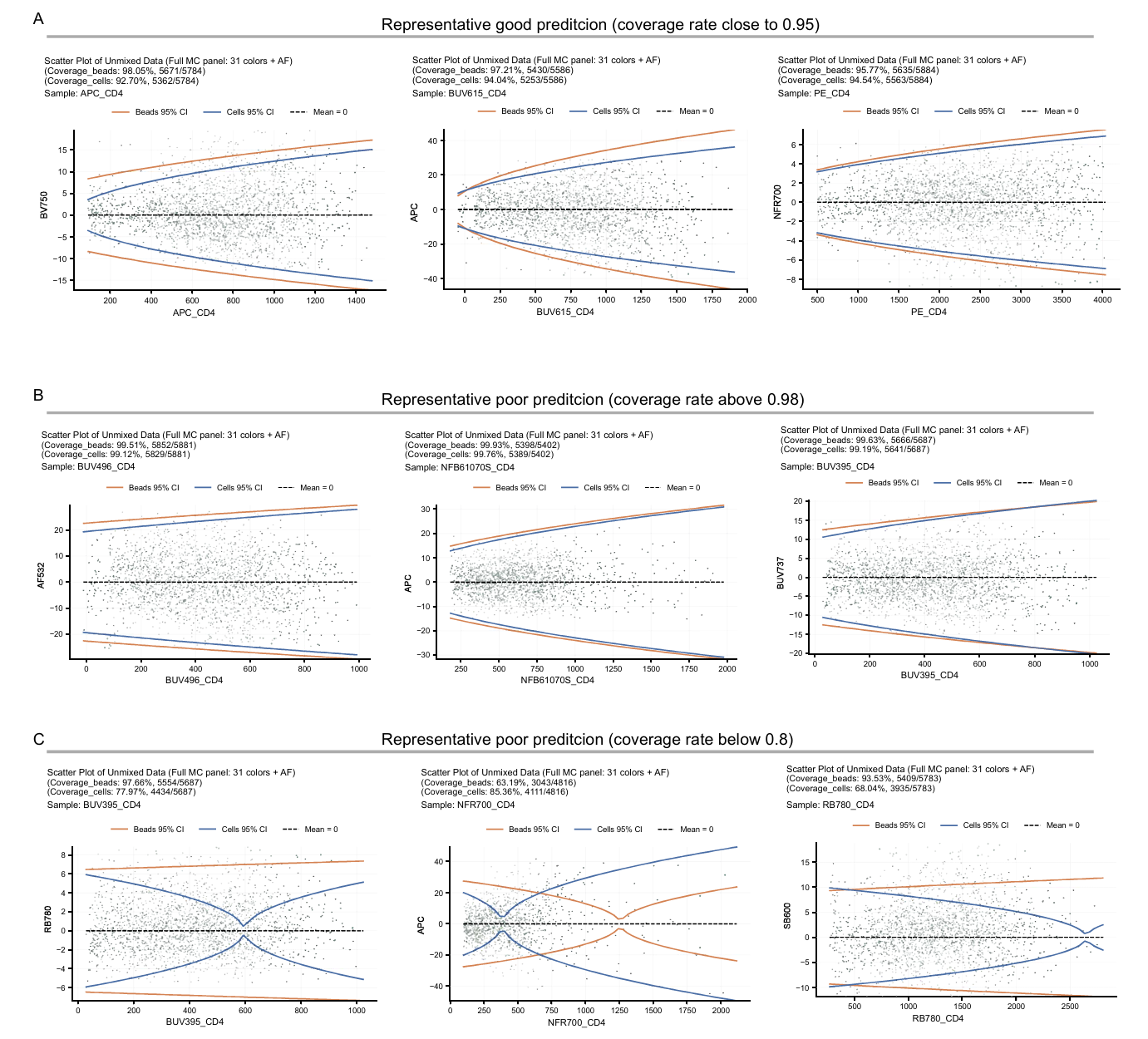

#### **Supplementary Figure 2 representative good and poor prediction of external validation.**

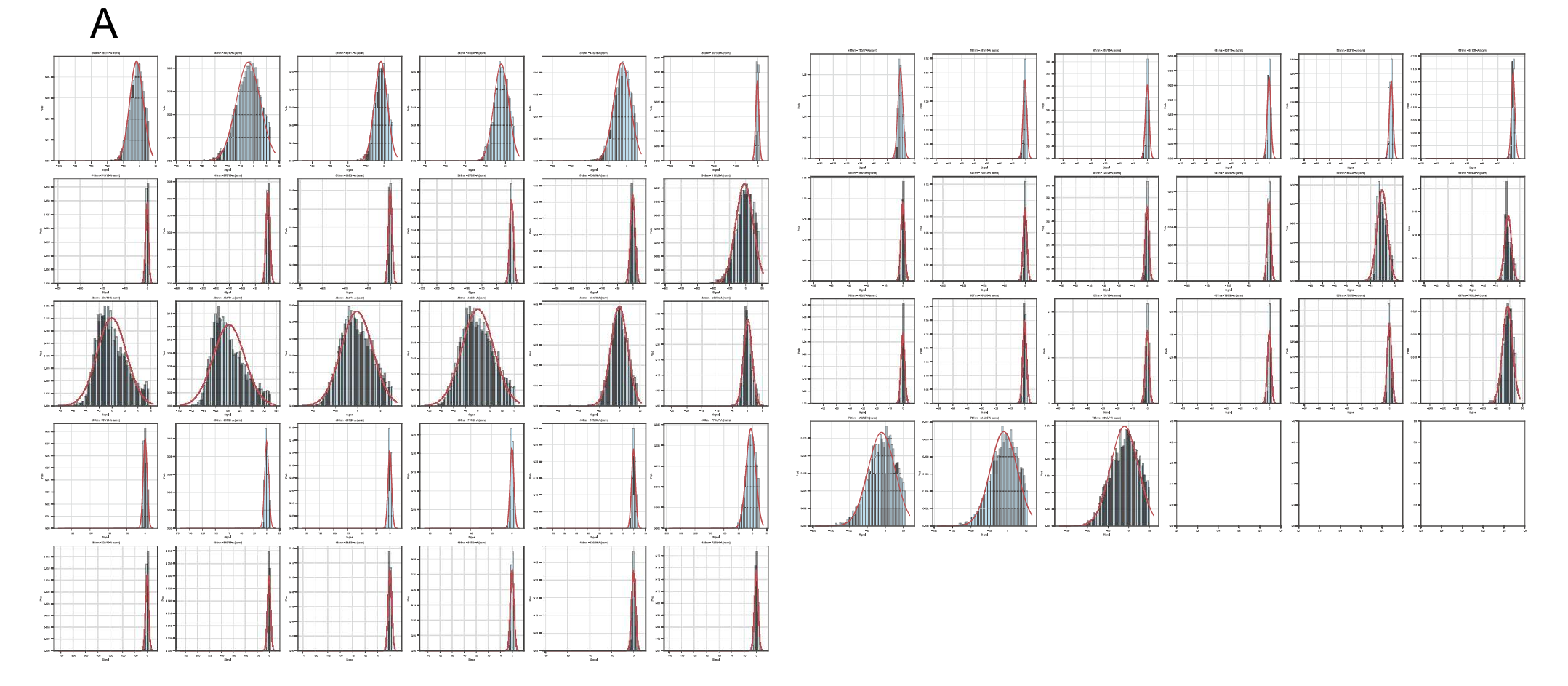

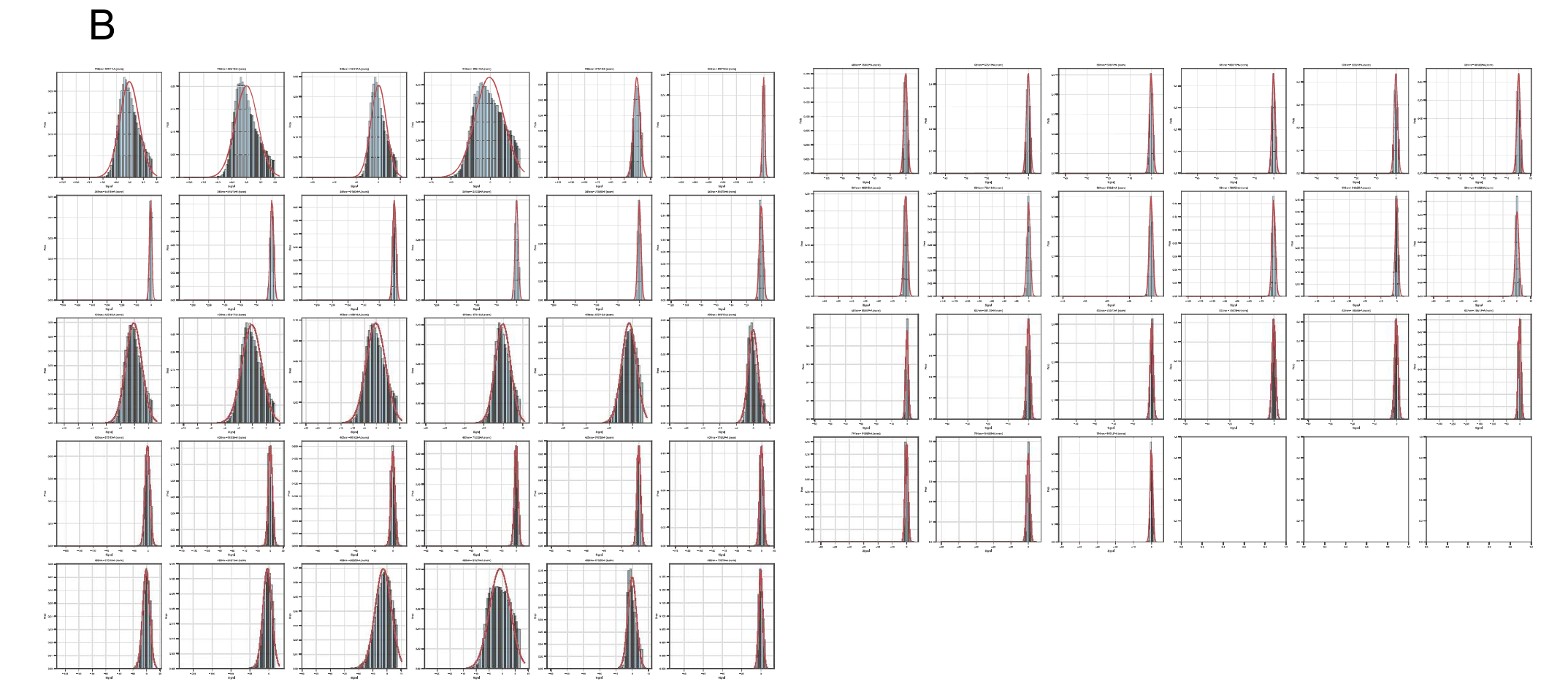

#### **Supplementary Fig3 the residual follows a normal distribution at each detector.**

(**A-B**) The distribution of residuals at each detector from the positive population of SCC cell BUV805-CD16(**A**) and FITC-CD2(**B**).

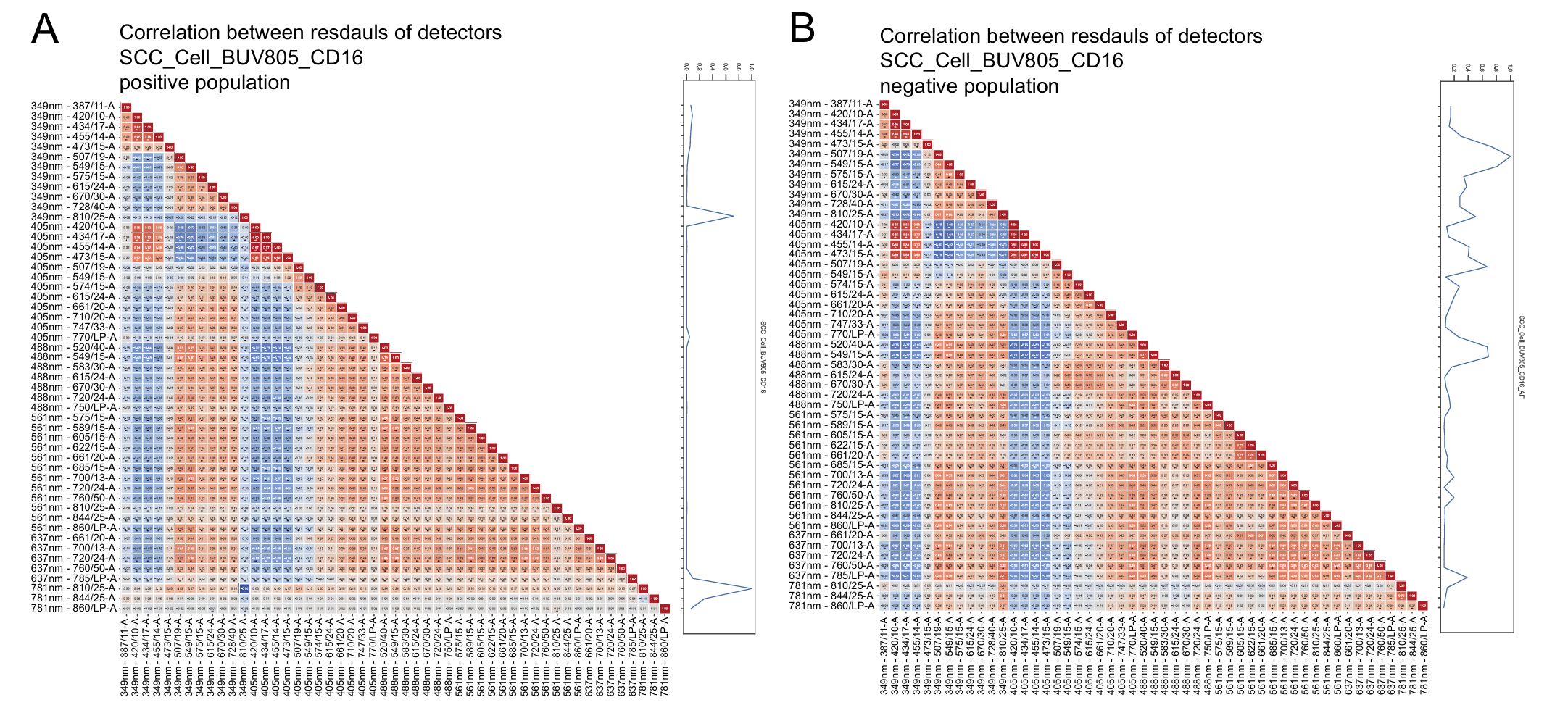

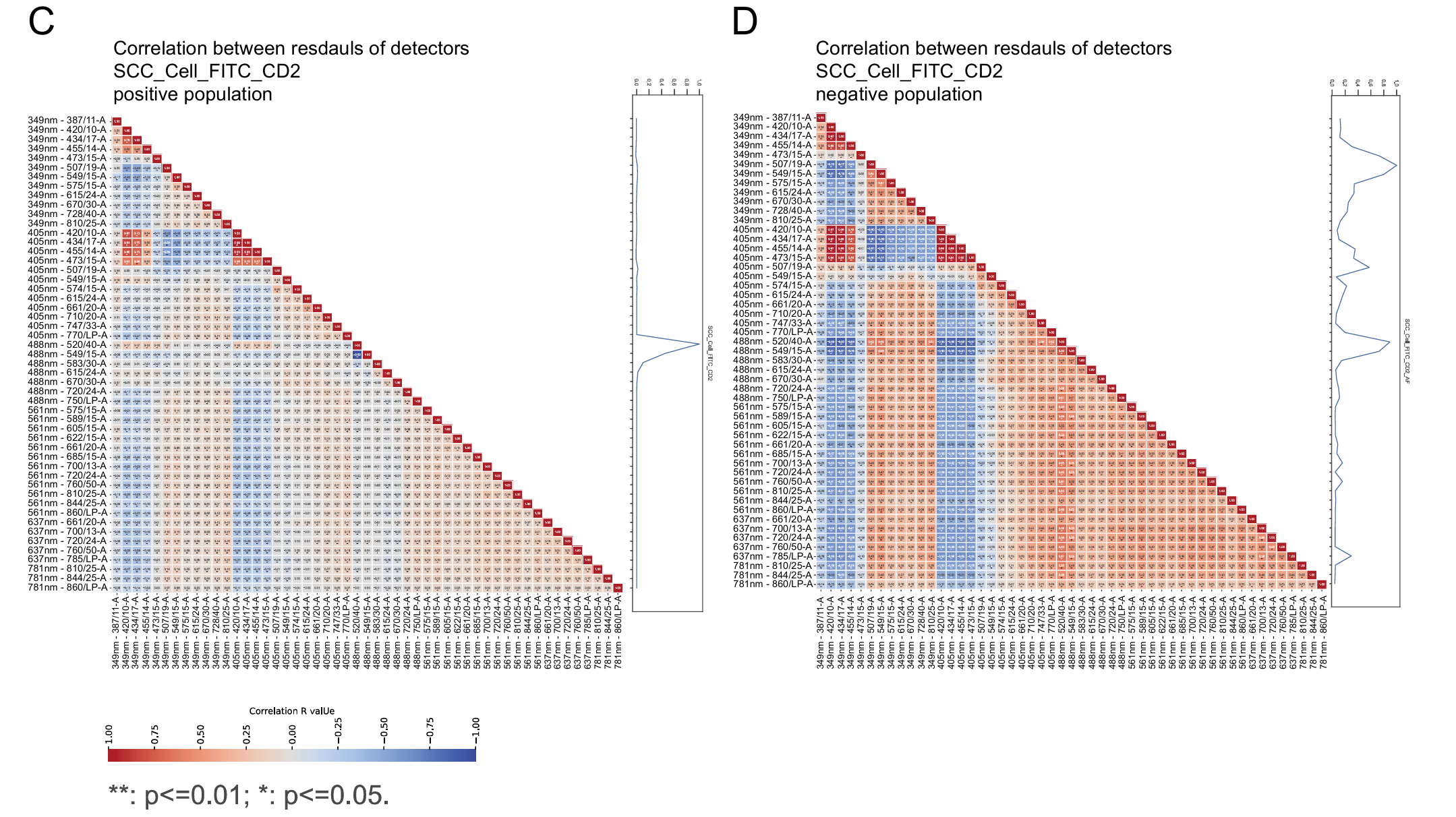

#### **Supplementary Fig4 The residuals at distinct detectors are correlated with each other**.

(**A-B**) The heatmap of correlation matrix of residuals at each detectors from the positive population (**A**) and negative population (**B**) of SCC cell BUV805-CD16 sample. (**C-D**) The heatmap of correlation matrix of residuals at each detectors from the positive population (**C**) and negative population (**D**) of SCC cell FITC-CD2 sample.

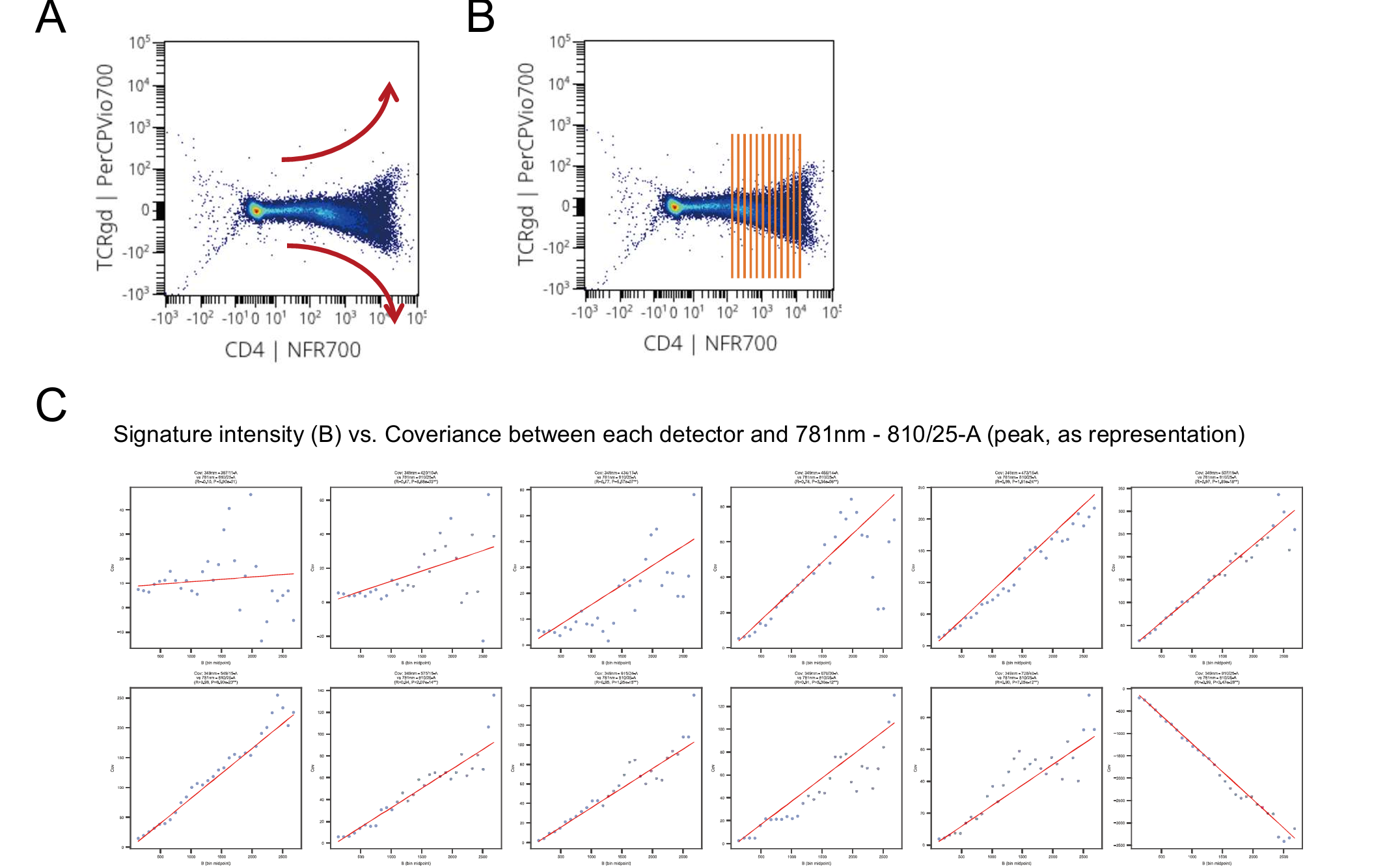

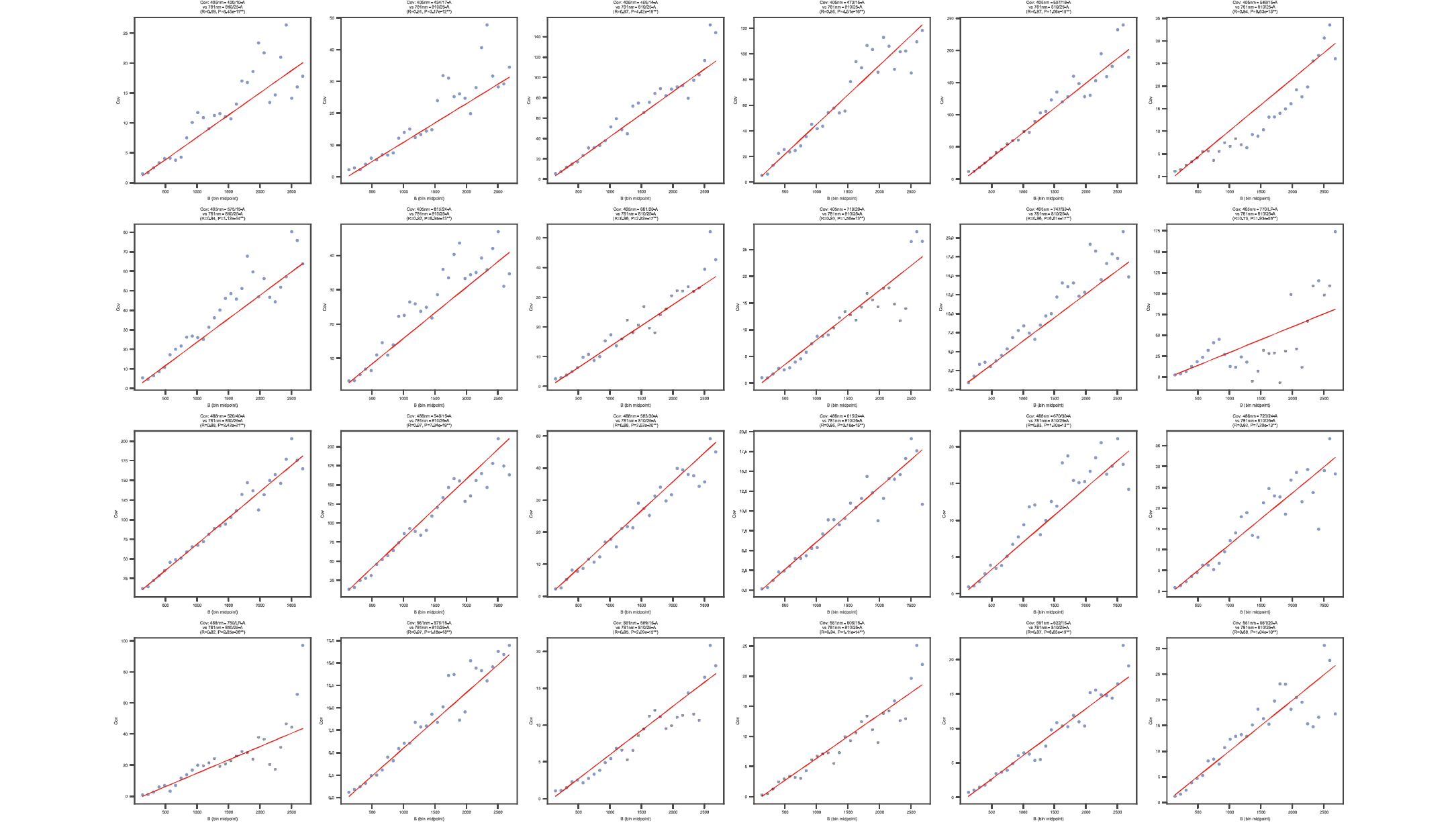

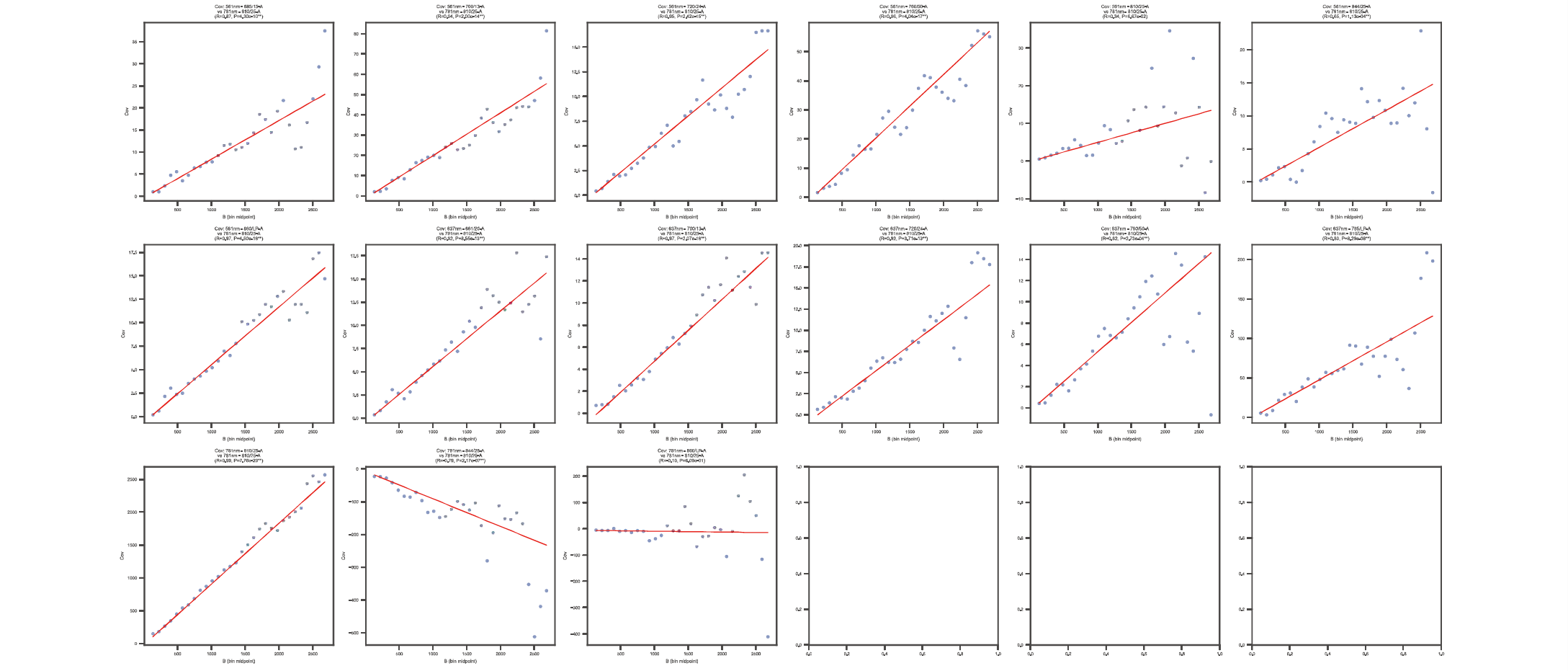

#### **Supplementary Fig5 The residuals are fluorescence intensity dependent.**

(**A**) example scatter plot shows the observed increase of spread along with the increase of fluorescence intensity. (**B**) Illustration of the subset method for the estimation of the observed increase of spread. (**C**) Example scatters of signature intensity (B) vs. coveriance between each detector and 781nm - 810/25-A (peak, as representation) from SCC cell BUV805-CD16 sample.
